# Developmental control of DNA damage responses in α- and β-cells shapes the selective beta-cell susceptibility in diabetes

**DOI:** 10.1101/2025.11.10.687707

**Authors:** Sneha S. Varghese, Alessandro Giovanni Hernandez-De La Peña, Aparamita Pandey, Laura Anchondo, Xiwei Wu, Supriyo Bhattacharya, Sangeeta Dhawan

**Affiliations:** Department of Translational Research and Cellular Therapeutics, Arthur Riggs Diabetes and Metabolism Research Institute, City of Hope, Duarte, CA 91010, USA; Biomedical and Biological Sciences Program, Keck School of Medicine, University of Southern California, Los Angeles, CA 90027, USA; Division of Pulmonary and Critical Care Medicine, David Geffen School of Medicine, University of California Los Angeles, 700 Tiverton Drive, Los Angeles, CA 90095, USA; Department of Molecular and Cellular Endocrinology, Arthur Riggs Diabetes and Metabolism Research Institute, City of Hope, Duarte, CA 91010, USA; Integrative Genomics Core, City of Hope, Duarte, CA 91010, USA

**Keywords:** DNA damage, endocrine differentiation, beta-cell, alpha-cell, epigenetics, Dnmt3a, DNA methylation

## Abstract

Accumulation of DNA damage drives β-cell dysfunction, senescence, and death in Type 1 and Type 2 diabetes. While α-cell dysfunction also contributes to disease pathology, they remain remarkably resistant to senescence and cell-death. The mechanisms underlying these differential responses to diabetogenic stress, particularly differences in their DNA damage vulnerability, remain unclear. We demonstrate that replication introduces a window of genomic vulnerability in both α- and β-cells during neonatal growth, with α-cells exhibiting higher replication rates and DNA damage. We show that neonatal β-cells resolve DNA damage more efficiently during mitosis and favor error-free repair, while α-cells compensate for their higher DNA damage vulnerability through increased cellular turnover. Using mouse models of overnutrition and diabetes, we show that β-cells exhibit greater vulnerability to terminal DNA damage and impaired repair capacity under diabetogenic stress, with compensatory replication amplifying this vulnerability. We demonstrate that developmental epigenetic programs shape the differential DNA damage vulnerability of postnatal β- and α-cells. Loss of *de novo* DNA methyltransferase Dnmt3a in pancreatic progenitors selectively increases the DNA damage vulnerability of β-cells from neonatal growth through adulthood. Our findings uncover novel developmental mechanisms that shape the distinct DNA damage responses of postnatal β- and α-cells during growth and diabetes.

**Article Highlights:**

- Mechanisms underlying the differential susceptibility of pancreatic β- and α-cells to diabetogenic stress remain unclear.
- Do replication dynamics, repair fidelity, and developmental epigenetic programs determine the vulnerability of postnatal β- and α-cells to DNA damage, a key driver of β-cell failure in diabetes?
- Replication introduces DNA damage vulnerability in both neonatal β- and α-cells, yet β-cells resolve damage more efficiently. Loss of DNA methyltransferase 3a in pancreatic progenitors selectively heightens β-cell DNA damage vulnerability that persists into adulthood. Moreover, diabetogenic-stress preferentially compromises β-cell repair fidelity.
- These findings reveal how developmental programs shape β-cell resilience and may influence lifelong diabetes risk.

## Introduction

Pancreatic islets maintain glucose homeostasis through the coordinated actions of the insulin-producing β-cells and glucagon-producing α-cells. Despite their shared developmental origin and proximity, these cells differ in their vulnerability to stress. In both Type 1 and Type 2 diabetes (T1D and T2D), progressive β-cell failure due to compromised identity, senescence, and cell-death impairs insulin secretion and drives hyperglycemia(1–5). While α-cells also become dysfunctional in diabetes, they remain resistant to destruction(6; 7). Understanding the molecular basis of this differential susceptibility could inform targeted therapeutic strategies to protect β-cell mass.

DNA damage is a key early catalyst underlying β-cell defects in all forms of diabetes(8–10). Failure to repair DNA breaks can compromise genomic integrity and cellular viability, a particular challenge for β-cells due to their long lifespan, limited self-renewal, and high propensity for endoplasmic reticulum (ER) stress and oxidative-stress(11). Combined with fluctuating metabolic demands, these factors heighten β-cells’ DNA damage burden. DNA damage response (DDR) pathways detect and repair DNA breaks, and their importance in β-cells is underscored by increased DNA breaks and DDR activation in both T1D and T2D, DNA repair polymorphisms linked to T1D, and diabetes onset in mice with β-cell repair defects(8–13).

Developmental trajectory further shapes cellular vulnerability. Establishment of mature β-cell pool requires tightly coordinated self-renewal and differentiation involving epigenetic programming and culminates in β-cells transitioning from a replicative, immature state to a post-mitotic, glucose-responsive state(14; 15). The replication and epigenetic remodeling required for self-renewal and transcriptional reprogramming during differentiation, are inherently prone to DNA breaks, with processes such as DNA methylation directing both lineage choices and genomic stability(16; 17). Immature, neonatal β-cells exhibit higher DNA damage compared to mature, adult β-cells(11). Whether this reflects ongoing replication and epigenetic programming and how it compares to α-cells, remains unknown.

In this manuscript, we compare the DNA damage vulnerabilities of β- and α-cells during neonatal development, metabolic-stress, and in murine models of T1D and T2D. Our data reveal that replication during neonatal growth confers DNA damage vulnerability to both lineages, with α-cells showing higher DNA damage corresponding to their higher replication rate. Notably, β-cells resolve DNA breaks more efficiently during G2/M and prioritize error-free DNA repair, while α-cells sustain higher DNA damage and death, with elevated turnover compensating for their vulnerability. We show that β-cells display higher DNA damage and poor repair compared to α-cells during metabolic-stress and diabetes, with compensatory replication contributing to β-cell vulnerability. Finally, we show that DNA methylation during pancreatic development shapes the differential DNA damage vulnerability of β- and α-cells. Loss of *de novo* DNA methyltransferase Dnmt3a in pancreatic progenitors selectively increases β-cell DNA damage vulnerability during neonatal growth, which persists into adulthood. Overall, these findings demonstrate how developmental events determine the long-term genomic resilience of β-cells and reveal a novel epigenetic mechanism underlying this process.

## Methods

### Animals

All animal experiments were performed according to protocols approved by the Institutional Animal Care and Use Committee (IACUC) at City of Hope. C57BL/6J animals were bred by mating males and females, and pups collected at different stages. The *db/db* (JAX 000642; BKS.Cg-*Dock7^m^*+/+*Lepr^db^*/J), C57BLKS/J (JAX 000662; BKS), NOD/ShiLtJ (JAX 001976), and NOR/LtJ (JAX 002050) strains were obtained from The Jackson Laboratory. BKS and NOR/LtJ mice served as controls for *db/db* and NOD/ShiLtJ mice. respectively. Mice were fed *ad libitum* and housed under a 12hr. light/dark cycle. Both sexes were used unless specified. Male mice were used for the HFD studies, as C57BL/6J females are protected against short-term HFD(18). We used female NOD mice as approximately 80% of females develop diabetes compared to only 20-40% of males(19). For the HFD studies, 1.5 months old male mice (n=8/group) were fed a control diet (10% KCal from fat; Research Diets D12450B), or HFD (55% Kcal from fat, Envigo TD.93075) for 1- and 8-weeks. NOD and NOR pancreata were collected at 4 and 8 weeks. *Dnmt3a^fl/fl^*and *Pdx1*-Cre mice(20; 21) were crossed to delete Dnmt3a in the pancreatic progenitor lineage (3aPKO), with a Cre-inducible Rosa26R-LSL-*YFP* reporter allele(22)incorporated for lineage tracing and monitoring Cre efficiency.

### Immunostaining, imaging, and morphometry

Immunofluorescence was performed on paraffin-embedded pancreatic sections using standard protocols from our previous studies(23; 24)(see also Supplementary methods). Primary antibodies were diluted in the blocking buffer (Supplementary Table 1). Donkey- and goat-derived secondary fluorescent antibodies (Jackson ImmunoResearch) were diluted 1:200. Nuclei were labeled with DAPI-containing antifade medium (Vector Labs). Cell death was detected by TUNEL assay (DeadEnd™ Fluorometric System, Promega) per manufacturer’s instructions. Imaging was done on a Leica DM6B microscope (Leica Microsystems) using the LAS X software or a Zeiss LSM 880/900 confocal microscope with AiryScan using ZEN software.

To quantify replication and DNA damage in β-cells, all islets were imaged and Insulin+ (Ins+), Ins+Ki67+, and Ins+ □H2AX+ cells was manually counted using ImageJ. Data was expressed as a percentage of Ins+ cells. DNA damage severity was assessed □H2AX pattern, with nuclear foci reflecting modest damage and pan-nuclear staining indicating severe, pre-apoptotic DNA damage(25). In 8-weeks-old NOD mice, □H2AX frequency was also quantified β- and α-cells for islets with and without immune infiltration. To quantify replication-associated β-cell DNA damage, Ins+Ki67+ □H2AX+ cells were counted and expressed as a percentage of (i) Ins+ cells (total β-cells with replication-associated damage), (ii) Ins+Ki67+ cells (fraction of replicating β-cells with DNA damage), or (iii) Ins+ □H2AX+ (fraction of damaged β-cells undergoing division). DNA damage during G2/M was evaluated using pHH3 (G2: punctate; M: solid nuclear staining). Ins+, Ins+G2+, Ins+M+, Ins+ □H2AX+, Ins+G2+ □H2AX+, and Ins+M+ □H2AX+ cells were counted and expressed as percentage of Ins+ cells to determine β-cells in G2 or M phases and as percentage of Ins+G2+, or Ins+M+ cells, for G2/M β-cells with DNA damage. Replication and DNA damage in α-cells were quantified similarly, using Glucagon. For cell-death analysis, Ins+, Glu+, and corresponding TUNEL+ cells were counted to determine the percentage of apoptotic β- and α-cells. At least 1500 endocrine cells (adult), 500 cells (P7), 700 cells (P21) were analyzed per sample, with equal number of Ins+ and Glu+ cells per comparison.

### Islet isolation

Murine islets were isolated as described(23), by pancreatic perfusion through the bile-duct with Liberase (Sigma-Aldrich). For neonatal isolations, pancreata from 5–8 pups were pooled and processed similarly without perfusion. Islets were hand-picked under a stereomicroscope and recovered overnight in RPMI 1640 medium (11 mM D-glucose, Invitrogen) supplemented with 100 IU/ml penicillin, 100 μg/ml streptomycin and 10% FBS (vol./vol.) at 37°C, with 5% CO_2_.

### Bulk RNA-sequencing

Total RNA was extracted from 30-50 hand-picked islets isolated from C57BL/6J mice at P5 (neonatal) and 2.5 months (adult) using the RNeasy Micro Kit (Qiagen). cDNA libraries were prepared from 250 ng total RNA using the KAPA HyperPrep Kit with RiboErase (HMR) (Kapa Biosystems) using manufacturer’s instructions. Libraries were validated using the Agilent Bioanalyzer and quantified with the Qubit High Sensitivity DNA Assay (Thermo Fisher Scientific). Libraries were sequenced on Illumina HiSeq2500 (2X 101 bp, paired-end mode, SBS v4 kits) and subsequent analysis was completed by the Integrative Genomics Core at City of Hope.

Paired-end RNA-Seq reads were processed by removing sequencing adapters and polyA tails, followed by alignment to the mouse genome (mm10) using STAR V2.6a(26). The gene-level read counts per sample were generated using HTSeq v0.11.1(27). Differential expression analysis was conducted using edgeR(28). Gene set enrichment analysis (GSEA) was conducted using the Gene Ontology Biological Process (GO:BP) dataset(29), employing the R package ClusterProfiler V4.0(30). See Supplementary methods for details.

### Single-cell RNA sequencing

Cells were captured using the Chromium X system and Next GEM Single Cell 3’ Reagent Kits v3. (10x Genomics), per manufacturer’s protocol, yielding ∼5,800 and ∼18,000 cells from the small and large neonatal islet samples respectively. Final sequencing libraries were assessed using the Agilent High Sensitivity DNA Chip and quantified with the Qubit High Sensitivity DNA Assay Kit (Thermo Fisher). Libraries were sequenced on an Illumina NovaSeq 6000 platform with the S4 Reagent Kit v1.5 (Illumina) at TGen, using a paired end read sequencing of 28 cycles for Read 1, 101 cycles for Read 2, and 10 cycles each for i7 and i5 index reads. Image analysis was performed with Real-time analysis (RTA) v3.4.4 software.

Raw fastq reads from the in-house neonatal murine samples were aligned to the mouse genome (mm10) followed by gene level quantification (CellRanger V7.0 (31)), filtering of doublets, ambient RNA correction and removal of low quality cells (Seurat V4.0(32)). The in-house and published adult murine datasets were integrated using Harmony V1.2.4(33), followed by normalization, UMAP projection, clustering (Seurat V4.0(32)) and cell type identification using known marker genes(34; 35) (Supplementary table 2). Normalized and integrated human islet single-cell data was obtained from HPAP PANC-DB(36). Cell cycle phases were identified using Tricycle(37). DDR pathway analysis was conducted using GSEA (ClusterProfiler V4.0(30)) and Geneset Variation Analysis (GSVA(38)) employing a curated set of non-redundant pathways from GO-BP(29). Differentially regulated pathways between two conditions were identified from the GSVA data using student’s *t*-test. See Supplementary methods for details.

### Statistics

All data were expressed as Mean±S.E. (standard error). Mean and SEM values were calculated from at least triplicates (biological) of a representative experiment. The statistical significance of differences was measured by unpaired Student’s *t*-test for experiments with two groups and a continuous outcome, while a two-way ANOVA with Šídák, Bonferroni, or Fisher’s LSD post-hoc tests was used for experiments with repeat measures. A *P* value < 0.05 indicated statistical significance. \**P*<0.05, \*\**P*<0.01, \*\*\**P*<0.005, \*\*\*\**P*<0.001.

### Data and Resources

All data are available in the article or supplementary materials. Datasets and resources are available from the corresponding author upon reasonable request. Sequencing datasets are in the process of being submitted to GEO and will be made publicly available upon publication.

## Results

### Replication introduces vulnerability to DNA damage in both neonatal α- and β-cells

As neonatal β-cells exhibit elevated DNA damage(39), we asked whether high replication rates during neonatal growth affect islet genomic integrity. We quantified DNA damage ( □H2AX) and replication (Ki67) in α- and β-cells of C57BL/6J mice at postnatal days 7 and 21 (P7 and P21, Fig. 1A, B; Supplementary Figs. 1A, B). Replication of both cell-types declined from P7 to P21, with α-cells displaying higher replication rates (Supplementary Figs. 1B, C). Concomitantly, DNA damage in both cell-types also declined with time, remaining higher in α-cells (Fig.1C).

**Figure 1.**
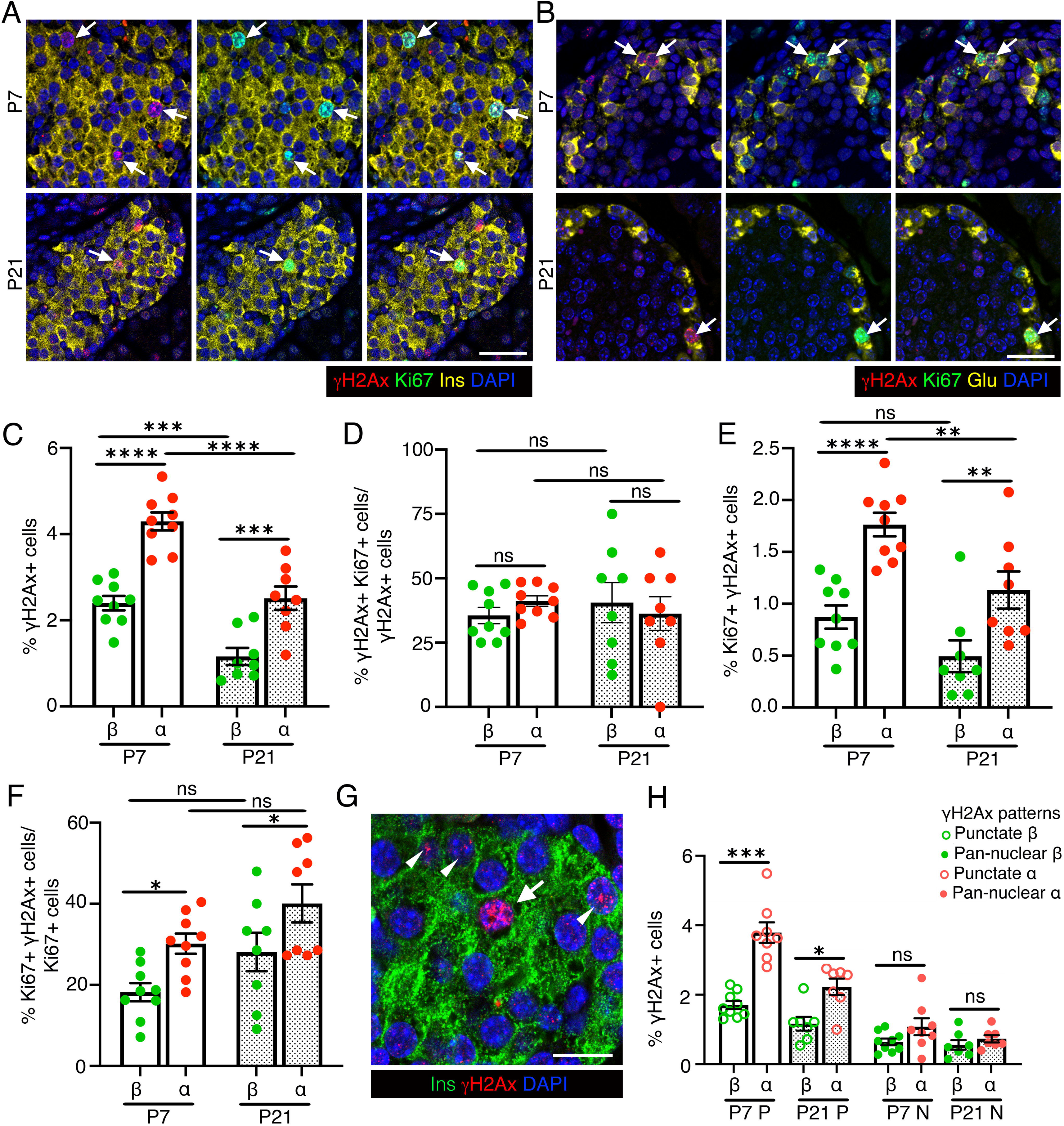
Replication introduces vulnerability to DNA damage in neonatal β- and α-cells. We analyzed and quantified DNA damage in the context of replication in β- and α-cells in mice at postnatal day 7 and 21, using □H2AX and Ki67 as respective markers. *(A, B)* Representative images showing immunostaining for □H2AX (red) and Ki67 (green) in β-cells *(A)* or α-cells *(B)*, with insulin (Ins) and Glucagon (Glu) shown in yellow, along with DAPI (blue) at postnatal day 7 and 21 (P7 and P21) after birth. Arrows indicate Ki67+ α- or β-cells marked by □H2AX. *(C)* Morphometric quantification of □H2AX+ β- and α-cells shown as percentage of total number of β- or α-cells at P7 and P21. *(D)* Quantification of □H2AX+Ki67+ β- and α-cells shown as percentage of □H2AX+ β- or α-cells at P7 and P21, highlighting the fraction of endocrine cells with DNA damage undergoing replication. *(E)* Quantification of Ki67+ □H2AX+ β- and α-cells shown as percentage of total β- or α-cells at P7 and P21, indicating the total pool of replicating endocrine cells with DNA damage. *(F)* Quantification of □H2AX+Ki67+ β- and α-cells shown as percentage of Ki67+ β- or α-cells at P7 and P21, showing fraction of the replicating endocrine cells harboring DNA damage. *(G)* Example immunostaining from a murine P7 pancreatic section showing □H2AX (red) in punctate (white arrowheads) and pan-nuclear (white arrow) in β-cells (Ins: green), with DAPI in blue. *(H)* Quantification of □H2AX+ β- and α-cells with punctate □H2AX+ (P) or pan-nuclear □H2AX+ (N) patterns shown as percentage of total number of β- or α-cells. All the images and quantification data shown is obtained from wildtype C57BL/6J pups at P7 (n=9 mice) and P21 (n=8 mice). *(C-F, H)* Red and green dots indicate α- and β-cell data points, respectively; white bars denote P7 data, dotted bars indicate P21 data. Error-bars show SEM. Ns= not significant, \**P<0.05,* ** *P*<0.01, \*\*\**P*<0.005, ****P<0.001, using 2-way ANOVA with Fisher’s LSD test for *(C-F)*, and paired, two-tailed *t*-test for *(H)*. Scale bar: 50 μm *(A, B)*, 20 μm *(G)*.

Quantification of DNA damage in replicating-cells showed that ∼50% of □H2AX + α- and β-cells at either stage were replicating, with no difference between lineages (Fig. 1D). The pool of replicating α- and β-cells with DNA damage declined from P7 to P21 as replication declined, remaining consistently higher in α-cells (Fig. 1E). Among replicating cells, a greater proportion of replicating α-cells harbored □H2AX foci than β-cells at both stages, the proportions remaining stable across development (Fig. 1F). α-cells more frequently displayed □H2AX punctae (moderate damage) than β-cells at both stages, while pan-nuclear □H2AX (pre-apoptotic damage) showed no difference but trended towards higher in α-cells (Fig. 1G, H). Overall, these data show that both replicating α- and β-cells are vulnerable to DNA damage, with higher α-cell replication rates corresponding to a larger fraction of cells with DNA breaks.

### α-cells have higher cellular turn-over and poorer capacity to resolve DNA breaks compared to β-cells during the growth phase

Effective DNA repair during replication is essential for cell-cycle completion and genomic integrity. To compare DNA repair efficiency in neonatal α- and β-cells during mitosis, we quantified □H2AX levels at P7 in cells marked by phosphorylated histone H3 (pHH3), distinguishing the G2 (speckled pHH3) and M (solid pHH3) phases (Fig. 2A, B. Supplementary Fig. 2A). Reflecting their higher replication, α-cells had more cells in G2 and M phases than β-cells, though their G2/M ratio was similar (Fig. 2C, Supplementary Fig. 2B). □H2AX patterns differed markedly between lineages during G2/M progression. Both G2 and M phase α-cells frequently harbored DNA damage, while β-cells showed minimal □H2AX in the M phase (Fig. 2D, Supplementary Fig. 2A). α-cells also displayed higher punctate □H2AX distribution (modest damage) in both phases and trended toward increased pan-nuclear labeling (severe damage) during M phase (Fig. 2E). These data suggest that β-cells resolve DNA breaks more efficiently before mitosis, whereas α-cells have limited repair capacity during replication. We hypothesized that α-cells compensate through higher turnover to eliminate damaged cells. TUNEL analysis at P7 confirmed higher α-cell death, consistent with increased turnover, a trend that persisted post-weaning but was not statistically significant (Fig. 2F, G, Supplementary Fig. 2C). Notably, α-cells maintained higher replication than β-cells during and after neonatal growth (Supplementary Figs. 1B, C). These findings reveal distinct strategies for maintaining genomic integrity during replication: β-cells efficiently repair DNA damage, while α-cells rely on increased turnover to offset poor repair capacity.

**Figure 2.**
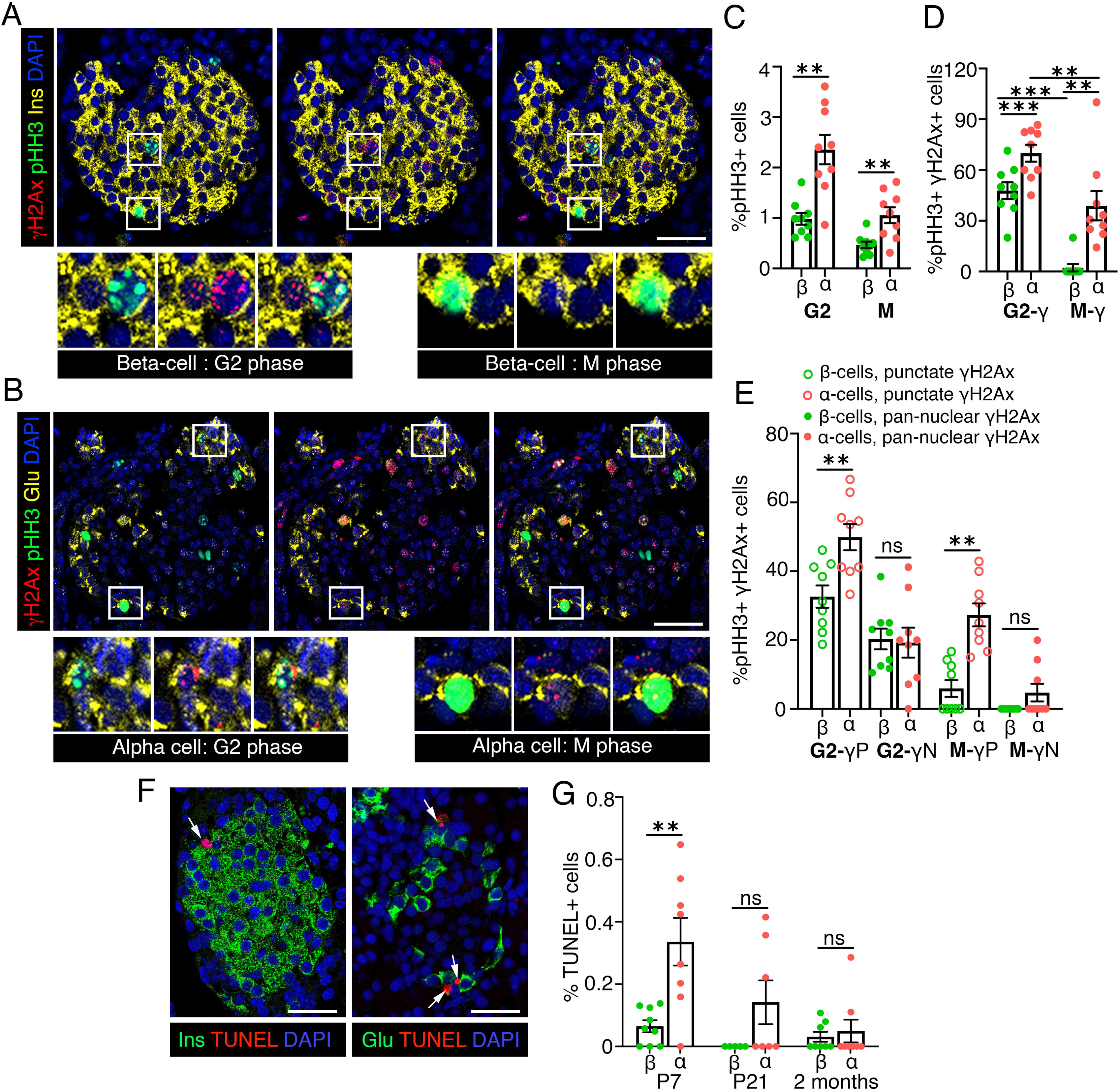
α-cells have poorer capacity to resolve DNA breaks and higher cellular turn-over compared to β-cells. We quantified DNA damage during the G2/M transition in β- and α-cells P7, using pHH3 to distinguish G2 and M phases. *(A, B)* Representative images showing Immunostaining for □H2AX (red) and pHH3 (green) in β-cells *(A)* or α-cells *(B)*, with insulin (Ins) and Glucagon (Glu) shown in yellow, along with DAPI (blue) at P7. Punctate pHH3 pattern indicates the G2 phase, while solid pHH3 staining represents the M phase. Regions marked by white squares in *(A, B)* indicate representative β- and α-cells in the G2 or M phases, shown below at 2X magnification. *(C, D)* Morphometric quantification of pHH3+ *(C)* and pHH3+ □H2AX+ *(D)* β- and α-cells, shown as percentage of total number of β- or α-cells at P7 *(C)* or as percentage of G2 or M phase β-cells *(D)*, respectively. *(D)* indicates the total pool of G2 or M phase endocrine cells with DNA damage. *(E)* Quantification of pHH3+ □H2AX+ β- and α-cells with punctate □H2AX+ (P) or pan-nuclear □H2AX+ (N) patterns shown as percentage of G2 or M phase β- or α-cells, to determine the fraction of the G2 or M phase endocrine cells with modest or severe DNA damage. *(F)* Representative images of a P7 murine pancreas showing TUNEL (red) in β- or α-cells (Ins/Glu: green), with DAPI in blue. *(G)* Quantification of TUNEL+ β- and α-cells at P7, P21, and 2 months of age. White arrows mark TUNEL+ endocrine cells. All the images and quantification data shown is obtained from wildtype C57BL/6J pups at P7 (n=9 mice), P21 (n=7 mice), and 2 months (n=8 mice). *(C, D, G)* Red and green dots indicate α- and β-cell data points, respectively. *(E)* Solid dots indicate pan-nuclear □H2AX+, while open dots indicate punctate □H2AX+ pattern. Error-bars show SEM. Ns= not significant, ** *P*<0.01, \*\*\**P*<0.005, using 2-way ANOVA with Fisher’s LSD test for *(D)*, paired *t*-tests for *(C, G).* For *(E)* paired two-tailed *t*-tests were used for both G2-P, G2-N, and M-P comparisons, while a Wilcoxson ranking test was used for M-N comparison. Scale bar: 50 μm.

### β- and α-cells employ distinct DDR programs during cell-cycle progression

To define the molecular basis of differential DNA damage vulnerability in α- and β-cells, we performed bulk RNA-seq on neonatal (P7) and adult (2.5-months old) C57BL/6J islets. Neonatal islets showed higher expression of replication and DDR genes and pathways and lower expression of β-cell maturation markers (Fig. 3A, B). Comparison of immature, stem-cell-derived and mature adult human-islet α- and β-cells using published scRNA-seq datasets (40; 41) confirmed the higher DNA damage vulnerability of immature endocrine cells in humans (Supplementary Figs. 3A, B). Single-cell RNA-seq (scRNA-seq) on neonatal (P7) C57BL/6J islets showed that Ki67+ cells were enriched in DDR pathways, including base excision and mismatch repair, confirming that replication is coupled with DDR activation (Fig. 3C, D, Supplementary Figs. 3C-F). To assess DDR dynamics during G2/M, we deduced cell-cycle phases using Tricycle, which assigns each cell a continuous theta (θ) value marking progression from early to late cell-cycle states. Tricycle assignments recapitulated canonical cell-cycle gene expression patterns, with *Ccne1/2* and *Mcm6* peaking at G1/S and *Cdk1* and *Aurka* at G2/M (Fig. 3E). Gene set variation analysis (GSVA) using GO-BP DDR terms identified 12 processes varying across G2/M in both cell-types and 14 unique to β-cells, indicating greater G2/M DDR remodeling in β-cells (Supplementary Fig. 3G). Among pathways differing between G2 and M phases, comparison of enrichment scores revealed stronger induction of alternative NHEJ, break-induced replication, inter-strand cross-link repair, and DNA gap filling in G2 β-cells (Fig. 3F). While error-free mechanisms such as DSB repair via HR remained active throughout G2 and M phases in both cell-types, error-prone translesion synthesis was downregulated in M phase β-cells but persisted in α-cells (Fig. 3F). These data suggest that β-cells enforce robust genomic quality-control via enhanced G2 repair and suppression of error-prone pathways in mitosis, while α-cells adopt more permissive repair corresponding to greater M-phase damage burden.

**Figure 3.**
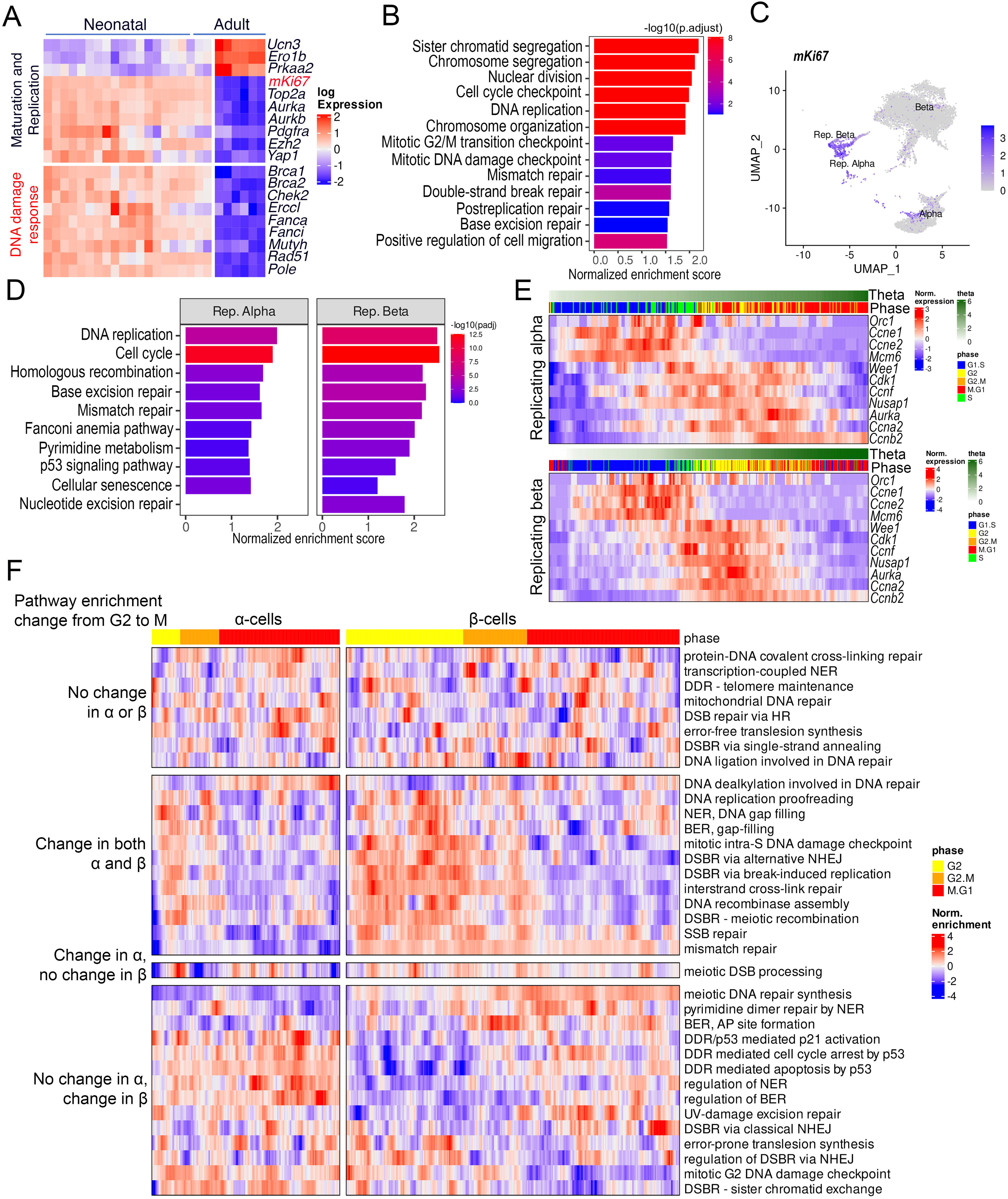
Comparison of DDR pathways among murine α- and β-cells during postnatal development and cell-cycle progression. DDR pathway enrichments were obtained by analyzing scRNA-seq and bulk RNA-seq profiles of neonatal and adult murine islets. Each heatmap row is normalized by converting into z-scores. In pathway enrichment bar-plots, bar heights and colors are proportional to the enrichment scores and adjusted *p*-values respectively. *(A)* Heatmap comparing the expressions of selected differentially expressed genes (*P*<0.05, replicates – neonatal: 20, adult: 6) associated with maturation/replication and DDR in neonatal and adult islet bulk RNA-seq. *(B)* Selected key replication/DDR pathways enriched in neonatal-compared to adult-islets bulk RNA-seq. Enrichments quantified through GSEA using the GO-BP database. *(C)* UMAP plot of single cell RNA-seq data from neonatal islets showing the locations of the replicative α- and β-cells, highlighted by the expression of the replication marker *Mki67*. *(D)* DDR pathways enriched in neonatal replicative α- and β-cells. Enrichments quantified through GSEA using the KEGG database. *(E)* Heatmaps showing the log-normalized expressions of key cell-cycle genes among replicative α- and β-cells from scRNA-seq of neonatal samples. Each heatmap column represents a single cell, ordered by their estimated θ parameter, signifying the cell-cycle stage. For clarity, the expression values are lightly smoothed by applying a convolution filter with a window width of 10 cells. The predicted cell-cycle phases are highlighted on top of each plot as colored horizontal bars. *(F)* Differentially enriched (*P*<0.05) DDR pathways (GSVA using the GO-BP database) between G2 and M phases compared among neonatal α- and β-cells. each column represents a single cell and enrichment values are smoothed as described in *(E)*. Pathways are categorized based on their comparative status between G2 vs M phases.

### Diet-induced metabolic stress selectively increases DNA damage vulnerability in β-cells

Overnutrition increases insulin demand, triggering β-cell hypersecretion and adaptive expansion by replication(42). The metabolic-stress due to increased β-cell workload and replication can induce DNA damage(9). To determine how overnutrition affects islet genomic stability, we quantified □H2AX in α- and β-cells in 8-weeks old, C57BL6/J mice fed with either control chow (CD) or a high fat diet (HFD) for 8 weeks (Fig. 4A). β-cells from HFD-fed mice showed higher DNA damage than CD controls, whereas α-cells showed no difference (Fig. 4B), suggesting that metabolic-stress preferentially impairs β-cell genomic integrity.

**Figure 4.**
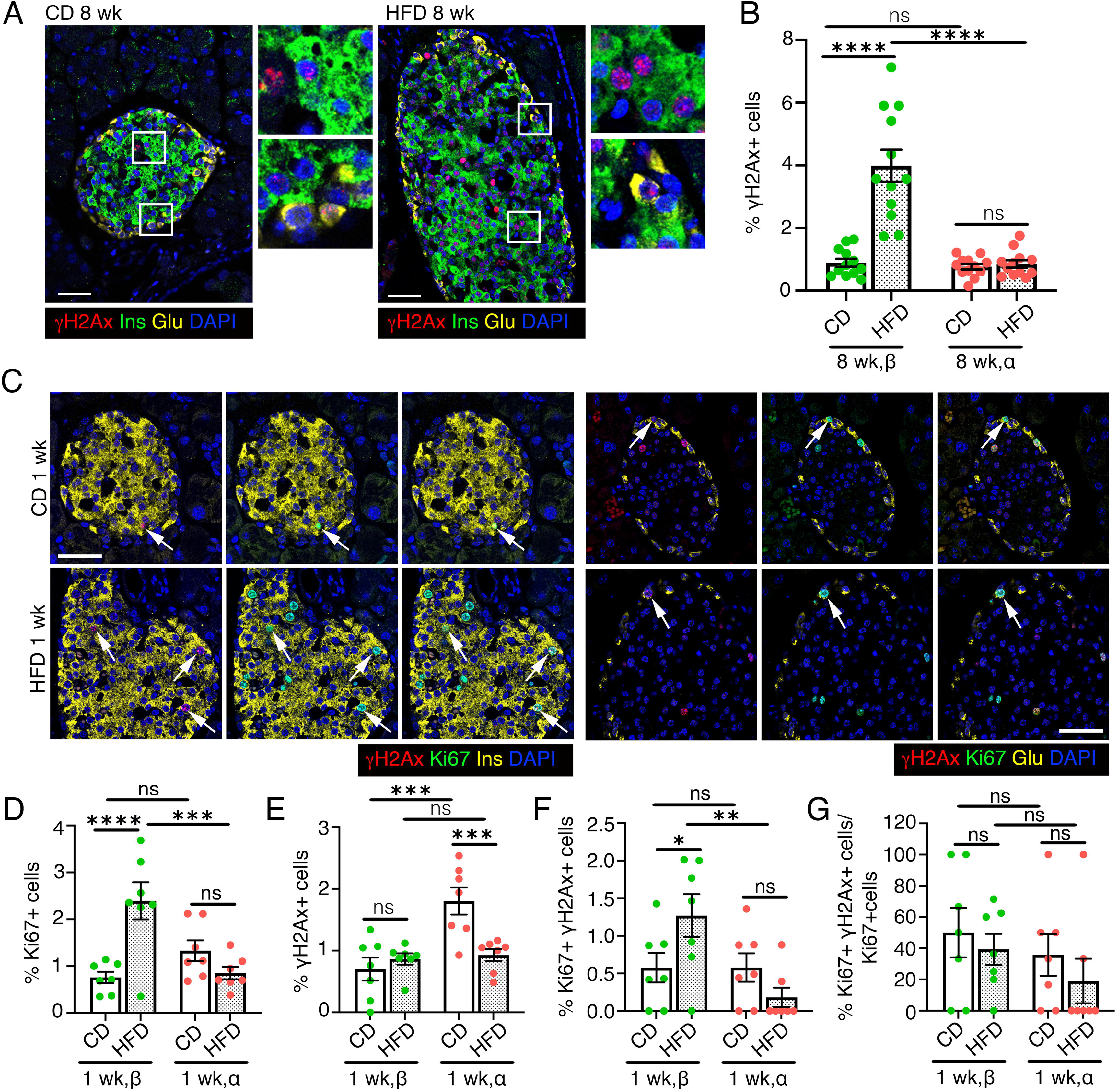
Metabolic stress increases the DNA damage vulnerability of β-cells. We quantified overall DNA damage as well as DNA damage in the context of replication in β- and α-cells in the high fat diet (HFD) induced metabolic stress model in mice with chow fed mice as a control. HFD administration was started at 6 weeks of age and continued for 1 or 8 weeks, with the control group remaining on chow diet (control, CD) throughout. *(A, B)* Representative images *(A)* and quantification *(B)* for □H2AX (red) in β-cells (left panels) or α-cells (right panels) in pancreatic sections from mice fed with a CD or HFD for 8 weeks, stained with insulin (Ins, green) and Glucagon (Glu, yellow), with DAPI (blue). Representative areas marked by white square corresponding to each image are presented as a 2X magnified panel next to the image. Data in *(B)* shows □H2AX quantification in β- and α-cells as percentage of total number of β- or α-cells in 8 weeks CD and HFD cohorts. *(C)* Representative images for □H2AX (red) and Ki67 (green) in β-cells (left panels) or α-cells (right panels), with insulin and Glucagon in yellow and DAPI (blue), in pancreatic sections from mice fed with CD or HFD for 1 week. Arrows indicate replicating α- or β-cells marked by □H2AX. *(D, E)* Quantification of Ki67+ *(D)* and □H2AX+ *(E)* in β- and α-cells shown as percentage of total number of β- or α-cells in CD and 1-week HFD cohorts. *(F, G)* Quantification of □H2AX+Ki67+ β- and α-cells shown as percentage of total *(F)* or Ki67+ *(G)* β- or α-cells in control and 1-week HFD mice. *(F)* indicates the total pool of replicating β- or α-cells with DNA damage, while *(G)* corresponds to the fraction of replicating β- or α-cells that harbor DNA damage. All the images and quantification data shown is obtained from wildtype C57BL/6J mice fed with an HFD or CD for 1 week (n=7 mice) or 8 weeks (n=12 mice), starting at 6 weeks of age. *(B, D-G)* Red and green dots indicate α- and β-cell data points, respectively; white bars denote control data, dotted bars indicate HFD data. Error-bars show SEM. Ns= not significant, \**P<0.05,* ** *P*<0.01, \*\*\**P*<0.005, ****P<0.001, using 2-way ANOVA with Fisher’s LSD test for *(B, D-G)*. Scale bar: 50 μm *(A, C)*.

To test if compensatory replication contributes to increased β-cell DNA damage, we quantified replication in both cell-types. β- or α-cell replication was similar between diets (Supplementary Figs. 4A, B), consistent with previous findings that β-cell replication peaks within first week of HFD treatment and declines thereafter(43). Thus, we compared DNA damage in the context of islet-cell replication after 1-week of HFD (Fig. 4C, Supplementary Fig. 4C). 1-week HFD increased β-cell, but not α-cell, replication (Fig. 4D) without affecting overall DNA damage (Fig. 4E). Correspondingly, more replicating β-cells carried DNA damage in HFD cohort compared to CD (Fig. 4F), whereas α-cells were not affected. While replicating HFD β-cells exhibited higher DNA damage than α-cells (Fig. 4F), the fraction of replicating β- and α-cells with DNA damage was similar between diets (Fig. 4G).

Analysis of □H2AX patterns revealed distinct temporal responses in HFD β- and α-cells. After 1 week of HFD, β-cells showed a trend towards higher punctate □H2AX with no difference in pan-nuclear □H2AX, while α-cells displayed reduced punctate and pan-nuclear labeling (Supplementary Fig. 4D). By 8 weeks, β-cells showed a significant increase in punctate □H2AX and no difference in pan-nuclear □H2AX, while α-cells remained unchanged (Supplementary Fig. 4E). TUNEL staining in 1- and 8-week cohorts showed increased β-cell death only after 8 weeks of HFD, with no difference in α-cell death at either time-point (Supplementary Figs. 4F-I). These data suggest that dietary stress induces progressive DNA damage in β-cells, driven in part by compensatory replication.

### β-cell vulnerability to DNA breaks is heightened in obesity associated Type 2 diabetes

Increased β-cell DNA damage is well documented in T2D and associated with β-cell senescence and failure. As hyperglycemia can induce DNA damage(9), we asked whether β-cells are more susceptible to DNA damage accumulation than α-cells in diabetes. Using *db/db* mice as a model of obesity-induced T2D along with background-matched BKS controls, we observed a β-cell-specific increase in □H2AX (both punctate and pan-nuclear) in 8-week-old *db/db* mice, a stage marked by hyperglycemia preceding diabetes onset (Fig. 5A-C, Supplementary Fig. 5A). α-cells showed no differences in DNA damage between groups. Given the compensatory islet expansion in *db/db* mice, replication (Ki67) was increased in both β- and α-cells (Supplementary Figs. 5B, C). The overall pool and fraction of replicating β- or α-cells with DNA damage did not differ significantly between strains, though *db/db* β-cells had a trend towards higher overall replication-associated damage (Fig. 5D, Supplementary Fig. 5D). Notably, we observed increased cell-death (TUNEL) in *db/db* β- but not α-cells (Fig. 5E, F). These data indicate that while both β- and α-cells undergo compensatory expansion in obesity-induced diabetes, only β-cells show increased DNA damage and cell-death, revealing a selective vulnerability that may reflect impaired DNA damage resolution.

**Figure 5.**
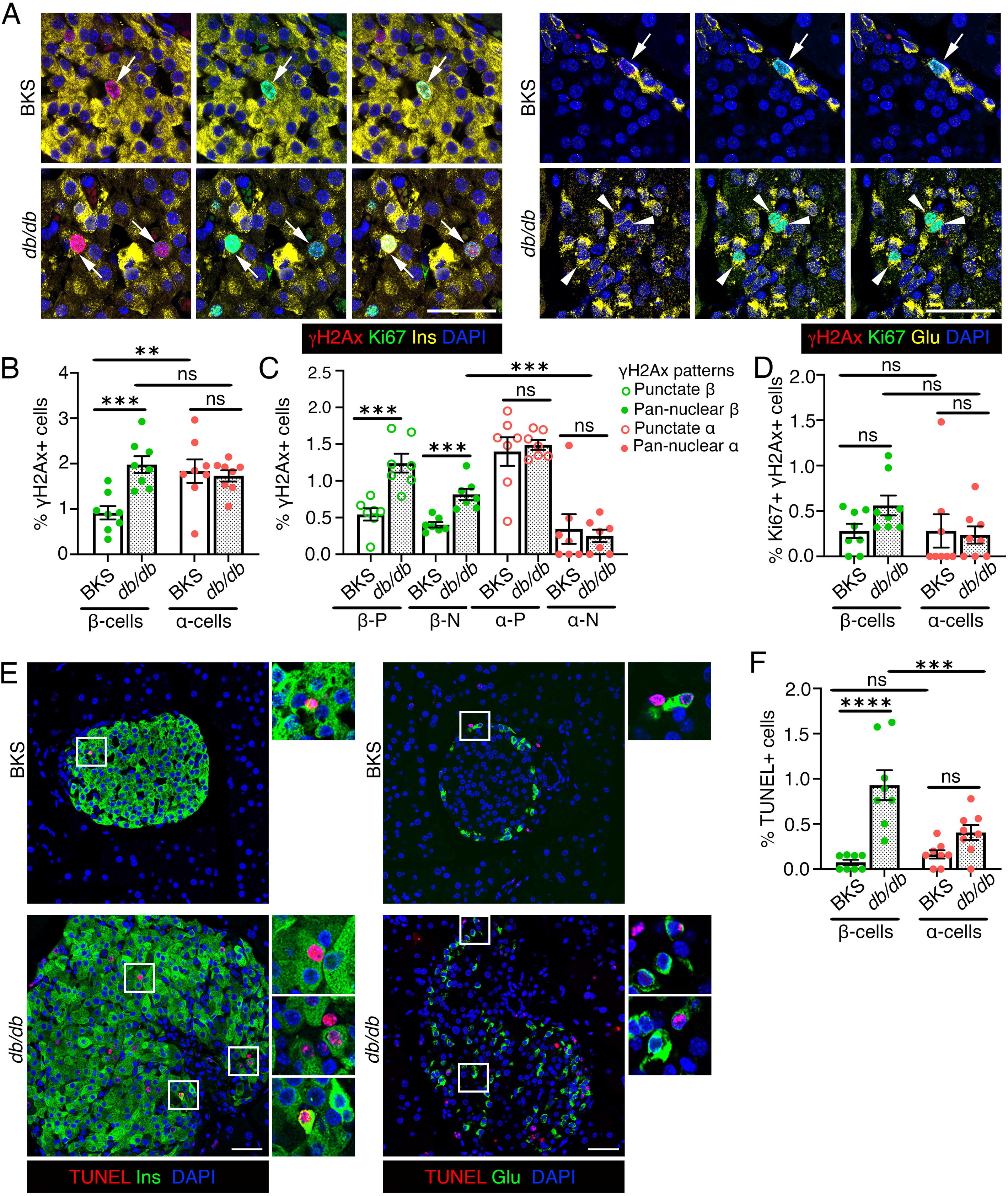
The susceptibility of β-cells to DNA damage is increased in an obesity-induced murine model of diabetes. We quantified DNA damage in the context of replication in β- and α-cells in the *db/db* mouse model of obesity induced diabetes and control, BKS mice, using □H2AX and Ki67 to detect DNA damage and replication. *(A)* Representative immunostaining images for □H2AX (red) and Ki67 (green) in β-cells (left panels) or α-cells (right panels), with insulin (Ins) and Glucagon (Glu) in yellow, with DAPI (blue) in pancreatic sections from 8 weeks old BKS and *db/db* mice. Arrows indicate Ki67+ α- or β-cells marked by □H2AX. Arrowheads indicate Ki67+ cells without any □H2AX. *(B, C)* Quantification of overall □H2AX+ (B), and punctate (P) □H2AX+ or pan-nuclear (N) □H2AX+ (C) in β- and α-cells shown as percentage of total number of β- or α-cells in BKS and *db/db* mice. *(D)* Quantification of Ki67+ □H2AX+ β- and α-cells as percentage of total β- or α-cells in *db/db* and control BKS mice, indicating the total pool of replicating cells with DNA damage. *(E, F)* Representative images (E) and quantification (F) of cell death using TUNEL (red) in β- or α-cells (Ins/Glu: green), with DAPI in blue in *db/db* and control BKS pancreata. Representative areas marked by white square corresponding to each image are presented as a 2X magnified panel next to the image. All the images and quantification data shown is obtained from 8 weeks old BKS and *db/db* mice (n=8/group), except panel C (n=7/group). *(B-D, F)* Red and green dots indicate α- and β-cell data points, respectively; white bars denote BKS data while dotted bars show *db/db* data. Error-bars show SEM. Ns= not significant, ** *P*<0.01, \*\*\**P*<0.005, ****P<0.001, using 2-way ANOVA with Fisher’s LSD test for *(B, D, F)*, and unpaired, two-tailed *t*-test for *(C)*. Scale bar: 50 μm *(A, E)*.

### Replication contributes to the increased DNA damage vulnerability of **β**-cells in early stages of autoimmune diabetes

Persistent DNA damage and increased DDR contribute to β-cell senescence and dysfunction in T1D(3; 8). While α-cells also become dysfunctional in T1D, it does not involve senescence(6; 7). To investigate cell-type-specific DDR differences in T1D, we quantified □H2AX in α- and β-cells in prediabetic NOD and control NOR female mice (4- and 8-weeks) (Fig. 6A, Supplementary Fig. 6A). At 4 weeks, both cell-types showed higher DNA damage in NOD mice than controls, with β-cells exhibiting greater DNA damage than α-cells (Fig. 6B). By 8 weeks, only NOD β-cells maintained elevated DNA damage (Fig. 6B). Replication (Ki67) was higher in β-but not α-cells of 4-weeks old NOD mice than NOR, β-cells within NOD islets showing higher replication than α-cells (Fig. 6C, Supplementary Fig. 6B). No replication differences were observed between strains for either cell-type at 8 weeks (Fig. 6C). More replicating β-cells had DNA damage in 4-weeks NOD mice compared to NOR controls and to α-cells within NOD islets, although the fraction of replicating β-cells with damage was similar between strains (Fig. 6D, Supplementary Fig. 6C).

**Figure 6.**
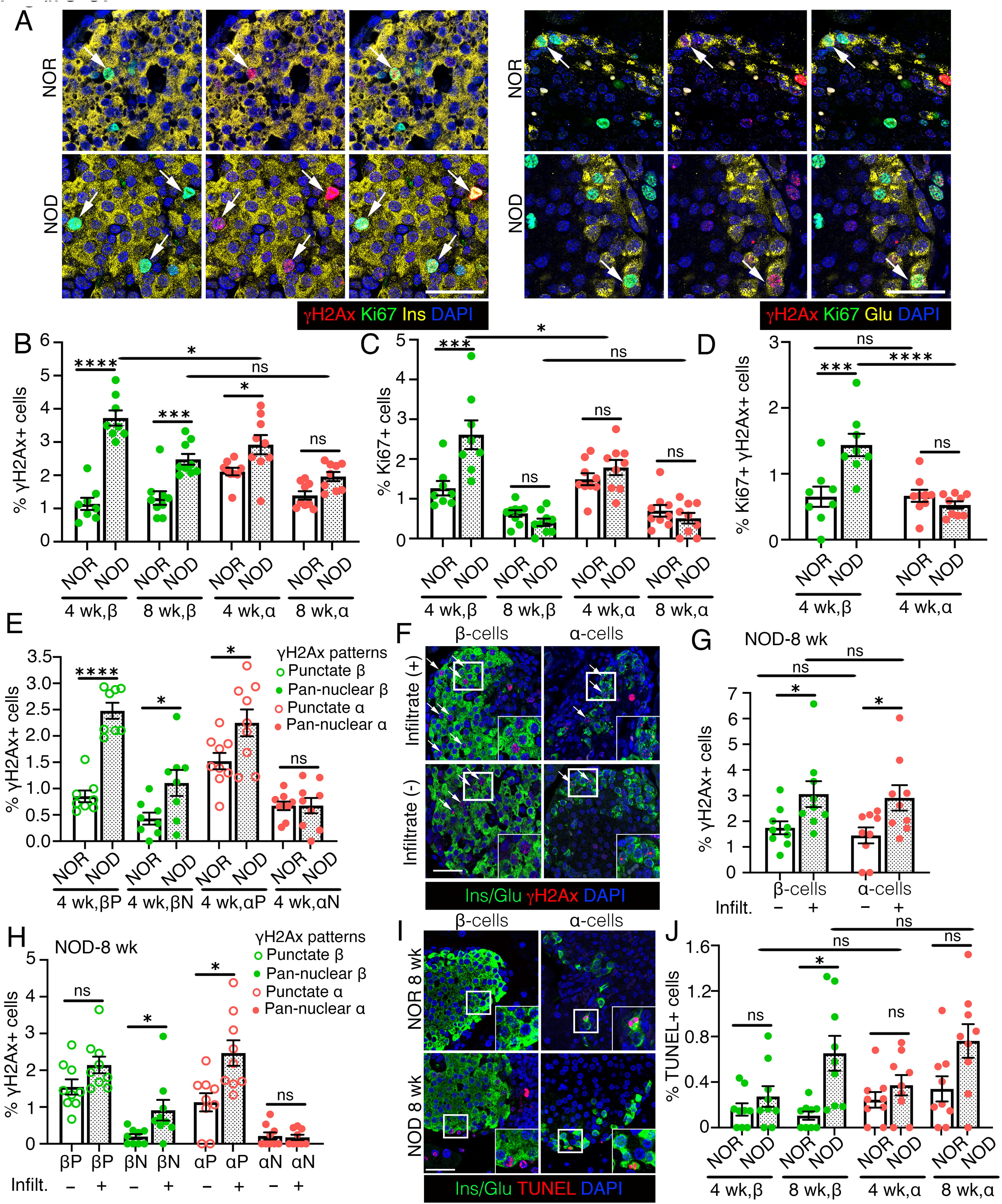
Replication contributes to the increased DNA damage vulnerability of β-cells during onset of autoimmunity in NOD mice. We quantified overall DNA damage as well as DNA damage in the context of replication in β- and α-cells in the NOD mouse model of autoimmune diabetes along with control NOR mice, using □H2AX and Ki67 to detect DNA damage and replication, respectively. *(A)* Representative images showing □H2AX (red) and Ki67 (green) immunostaining in β-cells (left panels) or α-cells (right panels), with insulin (Ins) and Glucagon (Glu) in yellow, with DAPI (blue) in pancreatic sections from 4 weeks old NOR (control) and NOD female mice. Arrows indicate Ki67+ α- or β-cells marked by □H2AX. *(B, C)* Quantification of □H2AX+ *(B)* and Ki67+ *(C)* β- and α-cells shown as percentage of total number of β- or α-cells in NOD and NOR mice at 4- and 8-weeks of age. *(D)* Quantification of □H2AX+Ki67+ β- and α-cells shown as percentage of total β- or α-cells in 4 weeks old NOD and NOR mice, indicating the total number of replicating cells with DNA damage. *(E)* Quantification of □H2AX+ β- and α-cells with punctate □H2AX+ (P) or pan-nuclear □H2AX+ (N) patterns shown as percentage of total number of β- or α-cells, in 4 weeks old NOD and NOR mice. *(F, G)* Representative images (F) and quantification (G) of □H2AX+ in β- and α-cells in islets with or without evident immune cell infiltrate, shown as percentage of total number of β- or α-cells in 8-weeks old NOD mice. □H2AX (red), Ins/Glu (green), DAPI (blue) in (F). *(H)* Quantification of □H2AX+ β- and α-cells with punctate □H2AX+ (P) or pan-nuclear □H2AX+ (N) in islets with or without infiltrate, shown as percentage of total number of β- or α-cells, in 8-weeks old NOD mice. *(I, J)* Representative images (I) and quantification (J) of cell-death using TUNEL (red) in β- or α-cells (Ins/Glu: green), with DAPI in blue, in 4- and 8-weeks old NOD and control NOR mice. The images in (I) are from 8-weeks old cohorts. All the images and quantification data shown is obtained from NOD and NOR mice at indicated ages (n=9/group). Representative areas marked by white square corresponding to each image are presented as a 2X magnified inset in (F) and (I). Red and green dots indicate α- and β-cell data points, respectively; white bars denote NOR data while dotted bars show NOD data. Ns= not significant, * *P*<0.05, ** *P*<0.01, \*\*\**P*<0.005, ****P<0.001, using 3-way ANOVA with a Tukey’s post-hoc test for *(B, C, J)*, 2-way ANOVA with Fisher’s LSD test for *(D, G)*, and paired, two-tailed *t*-tests for *(E, H)*. Scale bar: 50 μm *(A, F, I)*.

Analysis of DNA damage severity showed that both cell-types had increased punctate □H2AX (modest damage) in 4-weeks old NOD mice, but only β-cells showed increased pan-nuclear □H2AX (terminal damage) (Fig. 6E, Supplementary Fig. 6D). Punctate □H2AX remained higher in both cell-types at 8-weeks NOD, with no differences in terminal damage in either cell-type compared to NOR (Supplementary Fig. 6E). As β-cell DNA damage correlates with immune-infiltration(9), we compared α- and β-cells in islets with and without infiltrate in 8-weeks old NOD mice. Both cell-types showed increased □H2AX in the infiltrated islets, but only β-cells showed increased terminal DNA damage (Fig. 6F-H). Moreover, apoptosis was selectively increased in β-cells of 8-weeks old NOD mice (Fig. 6I, J). These data suggest that impaired DNA damage resolution during compensatory replication in juvenile NOD mice exacerbates β-cell DNA damage and vulnerability.

### Metabolic stress impairs β-cell repair programs in mice and humans

To understand DDR mechanisms in adult quiescent α- and β-cells under homeostasis and stress, we meta-analyzed published scRNA- and bulk-RNA-seq datasets(44–46). Non-dividing α-cells consistently displayed higher DDR than β-cells in chow-fed adult mice, in agreement with our data (Fig. 7A). While these α-β differences observed in chow-fed mice largely remained unchanged after 1- or 8-weeks of HFD (Fig. 7A, Supplementary Fig. 7A)(44; 45), HFD increased the fraction of non-dividing β-cells enriched for p53-mediated apoptotic DDR(Fig. 7A-C, Supplementary Figs. 7A-C), consistent with increased DNA damage (Fig. 4).

**Figure 7.**
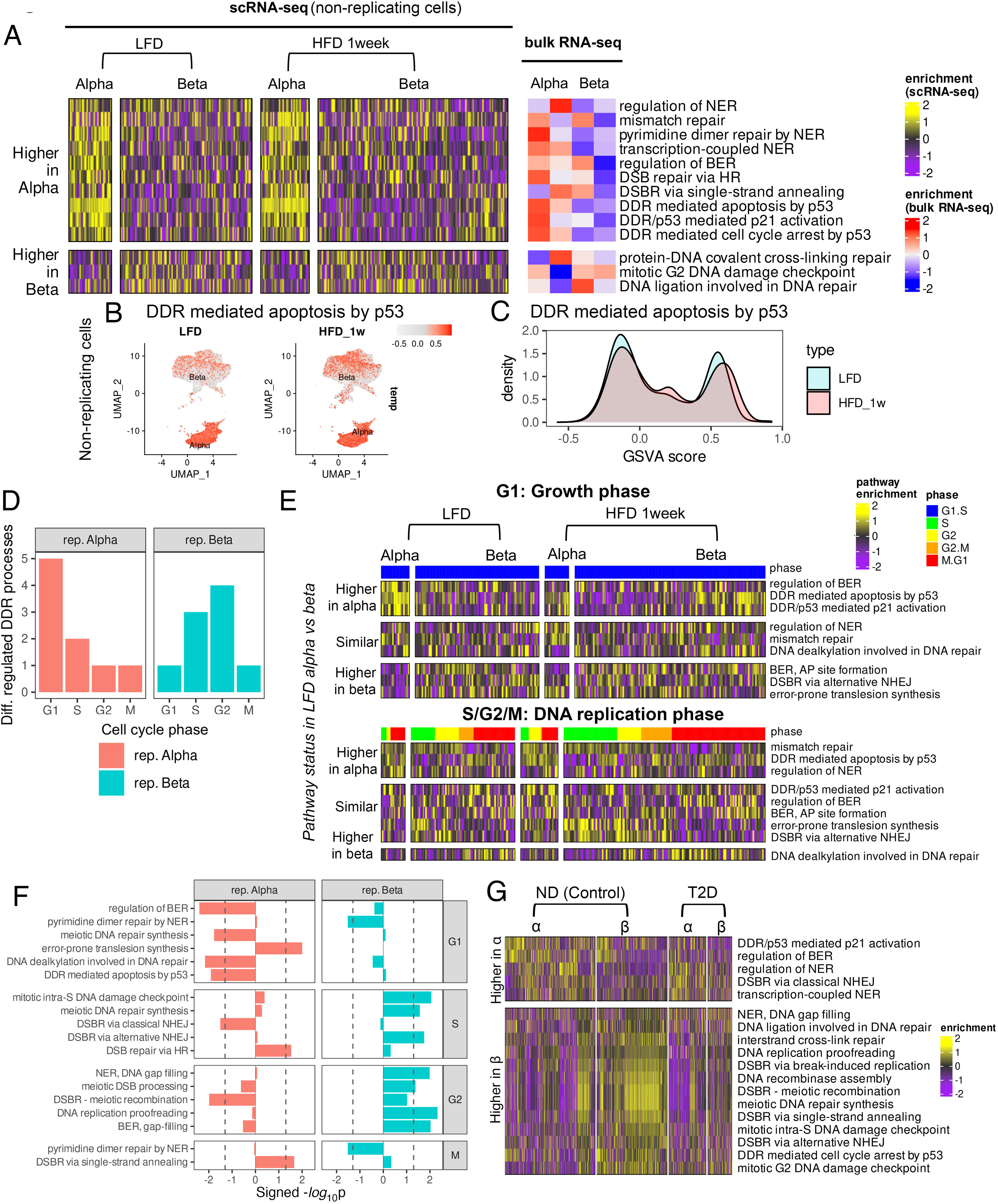
DDR differences among α- and β-cells under metabolic-stress in adult murine and human islets. HFD is used to model metabolic stress in mice. In humans, comparison between islets from non-diabetic and T2D donors was used to decipher the effect of metabolic stress. DDR pathway enrichments (GO-BP) are calculated using the GSVA method. *(A)* Heatmap showing differentially enriched (P<0.05) DDR pathways between adult murine non-replicative α- and β-cells under LFD and 1-week HFD from publicly available scRNA-seq data(44). Columns represent single-cell enrichment scores smoothed for clarity, as described in Fig. 3. For additional validation, GSVA enrichment scores from bulk RNA-seq of isolated α- and β-cells from an independent study are shown alongside. *(B, C)* UMAP plots and density distributions of enrichment score of the DDR mediated apoptosis pathway from scRNA-seq of murine non-replicative α- and β-cells compared between LFD and 1-week HFD. *(D)* GSVA scores of differentially enriched DDR processes between adult murine replicating α- and β-cells during different cell-cycle stages, compared between LFD and 1-week HFD. Heatmaps of DDR processes are shown separately for the G1 (initial growth) and S/G2/M (DNA replication) phases. Columns represent smoothed enrichment scores from single cells. *(E)* Number of differentially regulated DDR processes (P<0.05) between 1-week HFD and LFD as function of cell-cycle phase compared between α- and β-cells. *(F)* Differentially enriched DDR processes at each cell cycle phase. X-axis represents the magnitude of DDR difference between HFD, 1 week vs LFD, quantified as –log_10_(P). The sign indicates the direction of change. Positive values indicate higher enrichment in HFD, and vice versa. The dotted vertical lines represent – log_10_(0.05) as the threshold of significance. *(G)* Heatmap showing DDR process differences between human α- and β-cells from single cell RNA-seq of non-diabetic (control) and T2D donors (age: 30-50 years) from HPAP. Processes are categorized based on whether they are higher or lower in α-compared to β-cells among healthy donors.

To understand how metabolic-stress affects DNA repair during adaptive β-cell expansion, we compared DDR programs across cell-cycle phases in replicating α- and β-cells from 1-week chow (LFD) and HFD cohorts (Fig. 7D-F). HFD-induced DDR changes were most pronounced in G1 for α-cells and S/G2 for β-cells, with minimal changes in M phase (Fig. 7D), encompassing pathways both shared and distinct between LFD α- vs β-cells (Fig. 7E). In G1, HFD downregulated several DDR processes in α-cells, including base excision repair, DNA alkylation, and p53-mediated apoptosis (Fig. 7F). Despite minimal overall DDR remodeling in G1(Fig. 7F), HFD induced enrichment of p53-mediated apoptosis in a subset of G1 β-cells, similar to quiescent β-cells (Fig. 7E). In S/G2, β-cells upregulated the error-prone alternative NHEJ in response to HFD, while α-cells favored the accurate DSB repair via homologous recombination (HR)(Fig. 7E, F). This suggests that metabolic-stress compromises the robust repair that marks β-cell replication under homeostatic conditions. Comparison of human islet scRNA-seq data from non-diabetic donors (ND; 30-50 years, HPAP)(36) with murine LFD islet data revealed substantial cross-species conservation of α- vs β DDR differences (Supplementary Figs. 7D-F). Nine lineage-enriched DDR processes showed opposing α- vs β trends between species, with six enriched in mouse α-cells (*e.g.* p53 mediated cell-cycle arrest and mismatch repair) but higher in human β-cells (Supplementary Figs. 7D, E) likely reflecting their minimal replicative capacity. Notably, DDR processes typically enriched in ND β-cells were downregulated in donors with T2D (Fig. 7G), indicating impaired DNA repair consistent with high DNA damage.

### DNA methylation patterning by Dnmt3a during pancreatic progenitor differentiation directs the differential DNA damage vulnerability of islet α- and β-cells

DNA methyltransferase 3a (Dnmt3a) establishes DNA methylation patterns that define β-versus α-cell identity and β-cell maturation (24; 47; 48). As DNA methylation also maintains genomic stability(16; 17), we hypothesized that Dnmt3a-dependent DNA methylation during differentiation shapes DNA damage susceptibility of α- and β-cells. To test this, we assessed DNA damage and replication in neonatal (P7) α- and β-cells from mice lacking Dnmt3a in pancreatic-progenitor lineage (3aPKO) and littermate controls (Fig. 8A, Supplementary Fig. 8A-C). β-cells in 3aPKO mice showed significantly higher DNA damage than controls, while α-cells were unaffected (Fig. 8B). β-cell replication was reduced in 3aPKO mice, with no difference in overall pool of replicating α- and β-cells harboring DNA damage. Despite reduced replication, the fraction of replicating β-cells with DNA damage increased in 3aPKO mice, while α-cells showed no difference (Fig. 8C-E). Consistent with this selective vulnerability, 3aPKO β-cells also displayed increase punctate, but not pan-nuclear, □H2AX and increased apoptosis, compared to controls (Fig. 8F, Supplementary Fig. 8D-E).

**Figure 8.**
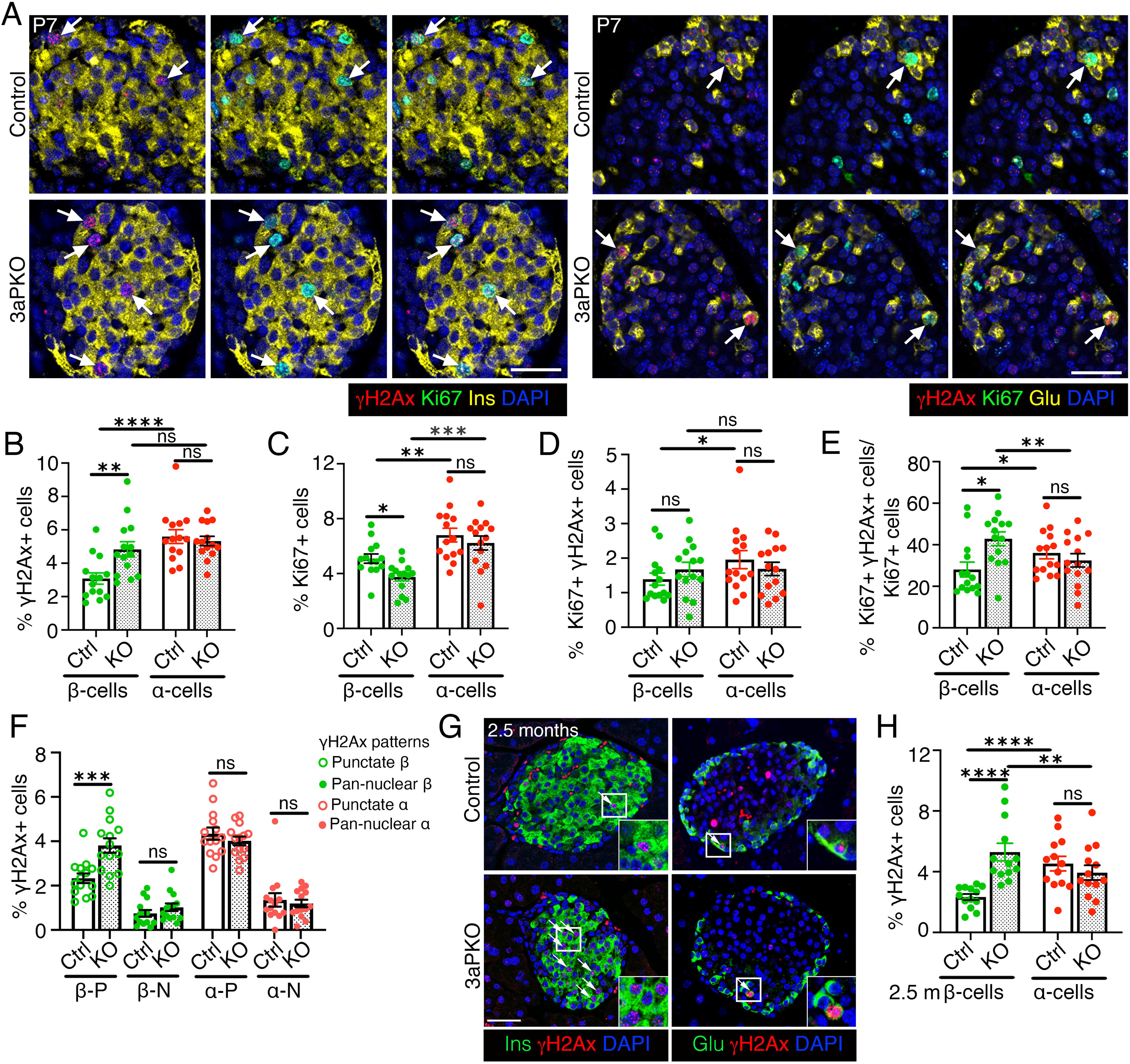
Dnmt3a action during pancreatic progenitor differentiation determines the differential DNA damage vulnerability of α- and β-cells. We analyzed and quantified DNA damage in the context of replication in β- and α-cells in control and 3aPKO (Dnmt3a KO in pancreatic progenitor lineage), using □H2AX and Ki67 as markers. *(A)* Representative immunostaining images for □H2AX (red) and Ki67 (green) in β-cells (left) or α-cells (right), with insulin (Ins) and Glucagon (Glu) shown in yellow, along with DAPI (blue) in control and 3aPKO mice at P7. Arrows indicate Ki67+ α- or β-cells marked by □H2AX. *(B)* Morphometric quantification of □H2AX+ β- and α-cells shown as percentage of total number of β- or α-cells in P7 control and 3aPKO mice. *(C)* Morphometric quantification of Ki67+ β- and α-cells shown as percentage of total number of β- or α-cells in 3aPKO and control mice at P7. *(D)* Quantification of Ki67+ □H2AX+ β- and α-cells in 3aPKO and control mice, shown as percentage of total β- or α-cells at P7, indicating the total pool of replicating cells with DNA damage. *(E)* Quantification of □H2AX+Ki67+ β- and α-cells shown as percentage of Ki67+ β- or α-cells in P7 control and 3aPKO pancreata, showing fraction of the replicating cells harboring DNA damage. *(F)* Quantification of □H2AX+ β- and α-cells with punctate □H2AX+ (P) or pan-nuclear □H2AX+ (N) patterns in P7 3aPKO and controls, shown as percentage of total number of β- or α-cells. *(G, H)* Representative images *(G)* for □H2AX+ and quantification *(H)* in β-cells and α-cells in pancreata from adult, 2.5 months old control and 3aPKO mice. In *(G) □H2AX*+ (red) is shown with Ins/Glu(green) and DAPI (blue), arrows highlighting α- or β-cells marked by □H2AX. Representative areas marked by white square corresponding to each image are presented as a 2X magnified inset. The graph in *(H)* shows □H2AX+ β- and α-cells as percentage of total number of β- or α-cells. All the images and quantification data shown is obtained from control and 3aPKO at P7 (n=14 mice) and 2.5 months (n=13 mice). *(B-F, H)* Red and green dots indicate α- and β-cell data points, respectively; white bars denote control data, dotted bars indicate 3aPKO data. Error-bars show SEM. Ns= not significant, \**P<0.05,* ** *P*<0.01, \*\*\**P*<0.005, ****P<0.001, using 2-way ANOVA with Fisher’s LSD test for *(B-E, H)*, and unpaired, two-tailed *t*-test for *(F)*. Scale bar: 50 μm *(A, G)*.

Next, we asked whether Dnmt3a depletion impairs β-cell DNA damage resolution during maturation. Elevated DNA damage persisted in β-cells of adult (2.5-months old) 3aPKO mice with no differences in control and KO α-cells (Fig. 8G-H), indicating sustained β-cell vulnerability from neonatal life into adulthood. To define the underlying trajectory, we examined islets P14, as β-cells mature and replication progressively decreases. At P14, 3aPKO β-cells had higher DNA damage, but showed no difference in replication, replication-associated DNA damage, or cell-death compared to controls (Supplementary Figs. 8F-L). β-cell mass remained similar between groups at all stages, with a trend towards reduction at P7 (Supplementary Fig. 8M). These findings suggest that replication-associated damage underlies selective β-cell vulnerability in Dnmt3a KO, its sequelae most pronounced during peak replication. Wildtype P7 islets had a higher proportion of Dnmt3a+ β-cells than α-cells, while both cell-types expressed Dnmt3a during late-embryonic stages (Supplementary Figs. 8C, N), suggesting that divergence in Dnmt3a expression between β- and α-cells emerges during neonatal life. Together, our data show that disruption of DNA methylation during progenitor differentiation selectively compromises β-cell genomic stability.

## Discussion

Our study reveals fundamental differences in how β- and α-cells manage genomic stability, offering insight into β-cell vulnerability in diabetes. We previously established that neonatal β-cells are more vulnerable to DNA damage than adult β-cells(39). Here, we find high DDR activity in neonatal islets, with both α- and β-cells exhibiting replication-associated DNA damage but employing distinct repair strategies. Neonatal α-cells display higher replication and greater DNA damage persisting through mitosis, whereas β-cells effectively resolve DNA damage before mitosis. α-cells compensate for higher DNA damage through increased turnover, suggesting cell-type-specific G2/M checkpoint mechanisms for genomic integrity. Single-cell analysis suggests that α-cells rely on error-prone repair during mitosis, while β-cells restrict such mechanisms to enforce stringent genomic quality control. These distinct strategies reflect their divergent postnatal trajectories, with β-cells transitioning to a post-mitotic state and α-cells retaining replicative capacity.

Elevated β-cell DNA damage is a shared hallmark and driver of β-cell defects in T1D, T2D, and monogenic diabetes(8–10). Across different models of β-cell failure, we observe a consistent pattern of heightened DNA damage vulnerability in β-compared to α-cells, driven in part by replication-associated DNA damage. While diabetogenic-stress induces compensatory β-cell expansion, our data show that it selectively impairs DNA repair in β-cells, especially within the replication pool, thereby amplifying their genomic fragility. This replication-associated vulnerability could extend across species, as attempted β-cell replication has also been reported in humans with T1D. Single-cell analyses of mouse HFD and human T2D islets further confirm stress-induced impairment of β-cell DNA repair and increased DNA damage. These observations underscore fundamental differences in DNA damage resolution between β- and α-cells and suggest that metabolic-stress impairs the repair pathways safeguarding replicating β-cells. This replication-associated DNA damage vulnerability may explain the peak incidence of β-cell autoimmunity during infancy and puberty(49; 50), periods of high β-cell growth and turnover. Unrepaired DNA damage during this window can induce senescence, mutations, and heightened immunogenicity(11). Combined with genetic and environmental triggers, this may elevate T1D risk, while under metabolic-stress, it can drive β-cell dedifferentiation and dysfunction, predisposing to T2D.

Our findings reveal lineage-specific requirements for DNA methylation in maintaining genomic stability, underscored by the selective DNA damage vulnerability of neonatal β-cells in 3aPKO mice, despite Dnmt3a loss in both lineages. β-cells maintain Dnmt3a expression through postnatal growth, whereas α-cells downregulate Dnmt3a earlier, reflecting their distinct DNA methylation requirements. The heightened sensitivity of β-cells to DNA methylation perturbations supports the model that α-cells represent a default endocrine cell-fate(51), while β-cell identity requires continuous epigenetic reinforcement. β-cell differentiation may require more extensive chromatin remodeling, creating regions of genomic instability that require proper DNA methylation for protection. This potentially explains why α-cells, despite higher replication rates that should warrant robust DNA methylation maintenance, are less dependent on DNA methylation for genomic stability.

These data reinforce that DNA methylation patterns established during early pancreas development shape postnatal islet homeostasis(24). The 3aPKO mice exhibit a selective, sustained increase in β-cell DNA damage from neonatal growth to adulthood. At P7, 3aPKO β-cells show increased DNA damage and apoptosis despite reduced replication, suggesting that Dnmt3a is crucial for β-cell genomic integrity during rapid postnatal expansion. The elevated β-cell DNA damage persists in adult 3aPKO mice without increased cell-death, which alongside previously reported functional defects, suggests impaired adaptability to metabolic-stress. Conditions warranting increased β-cell replication may unmask such latent vulnerabilities, with long-term consequences. Of note, the DNA methylation patterns associated with β-cell identity, function, and survival are erased in T2D human islets(52; 53). Beyond DNA methylation, other epigenetic regulators such as YY1 and Cohesin are also critical for postnatal β-cell genomic stability, and their dysregulation in diabetes underscores the broader epigenetic vulnerabilities underlying β-cell failure(54; 55). Together, our findings reveal that replication dynamics, metabolic-stress, and epigenetic safeguards shape differential DNA damage vulnerability of α- and β-cells, providing mechanistic insight into heightened β-cell vulnerability in early life. Our data establish a framework for understanding how developmental programming intersects with genetic factors and environmental challenges to drive β-cell failure and influence diabetes risk.

## Article Information

## Acknowledgements

We thank Dr. Nazia Parveen, Victor Ruiz, Daniel Nam, Alexander Ham, and Salma Elliessy (Dhawan Lab) for technical support. We would like to thank Drs. Rupangi Vasavada, Hung-Ping Shih, and Adolfo Garcia-Ocaña (City of Hope) for helpful discussions. We are grateful to City of Hope shared resources and core facilities and thank Dr. Brian Armstrong (Light Microscopy Core) and Drs. Hanjun Qin, Jinhui Wang, and Min-Hsuan Chen (Integrative Genomics Core) for their guidance and support. This manuscript used data acquired from the database (https://hpap.pmacs.upenn.edu/) of the Human Pancreas Analysis Program (HPAP; RRID:SCR_016202; PMID: <u>31127054</u>; PMID: <u>36206763</u>). HPAP is part of a Human Islet Research Network (RRID:SCR_014393) consortium (UC4-DK112217, U01-DK123594, UC4-DK112232, and U01-DK123716).

## Author Contributions

S.D., S.S.V., and S.B. conceived and planned the study. S.S.V., A.G.H., A.P., and L.A. performed the experiments. S.S.V., A.G.H., A.P., L.A., X.W., S.B., and S.D. performed the analyses. S.S.V., A.G.H., S.B., and S.D. interpreted data. S.B. and S.D. wrote the original draft of the manuscript, S.S.V., A.G.H., S.B., and S.D. revised and edited the manuscript. S.S.V. and S.D. acquired funding.

## Guarantor Data Access and Responsibility Statement

S.D. is the guarantor of this work and, as such, had full access to all the data in the study and takes responsibility for the integrity of the data and the accuracy of the data analysis.

## Prior Presentation

A small part of this work has been presented previously as a poster at the American Diabetes Association 84^th^ Scientific Sessions, 2024.

