## Supplementary Fig, Supplementary Table for "Developmental control of DNA damage responses in α- and β-cells shapes the selective beta-cell susceptibility in diabetes"

**Supplementary Figures and Legends**

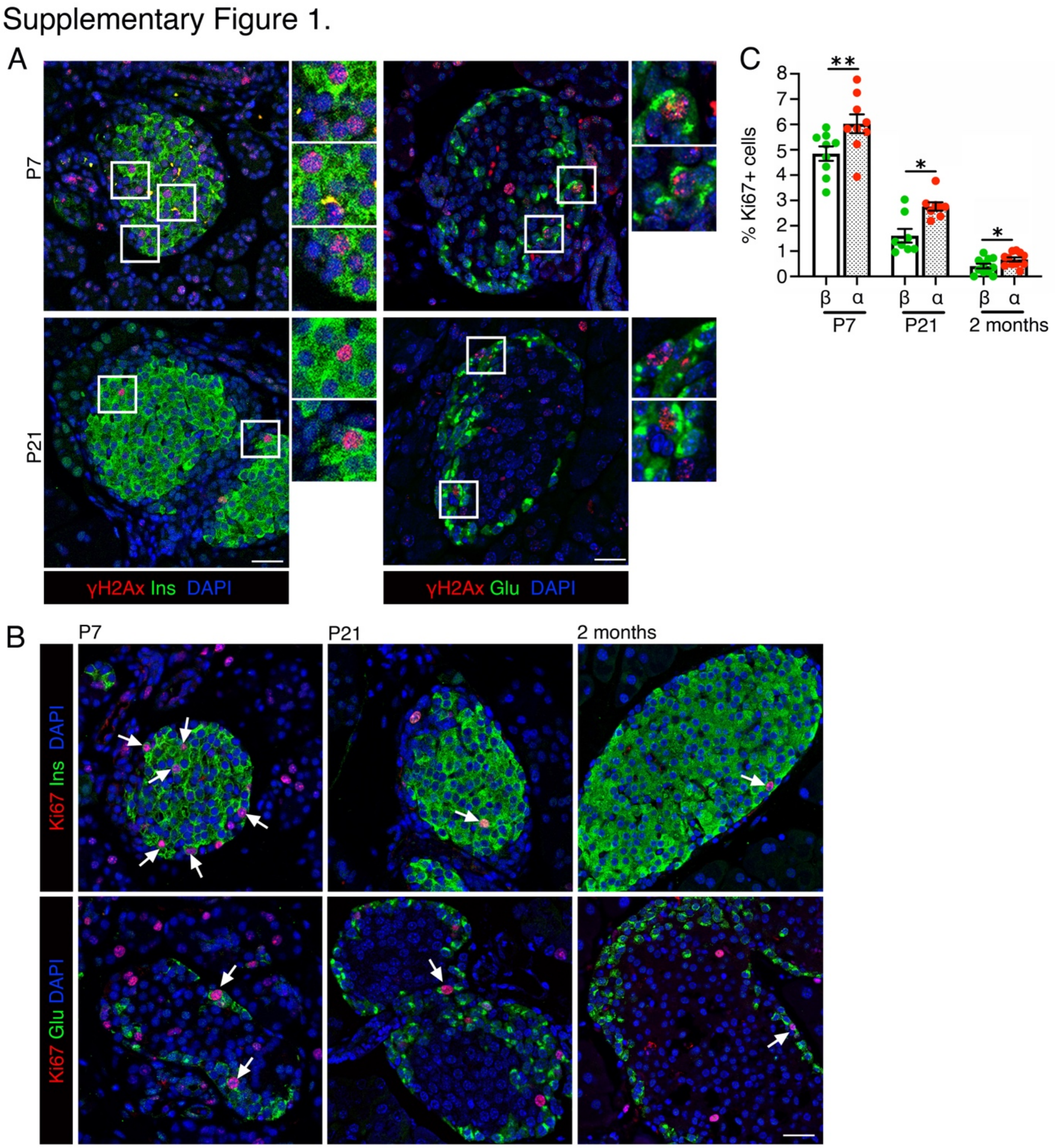


**Supplementary Figure 1 (Related to Figure 1).** *(A)* Representative images showing immunostaining for ɣH2AX (red) and β- or α-cells marked by insulin or glucagon (Ins/Glu: green), along with DAPI (blue) at P7 and P21. Representative areas marked by white square for each image are presented as an accompanying 2X magnification next to the image. *(B)* Representative images showing Immunostaining for Ki67 (red) in β- or α-cells labeled with insulin or glucagon (Ins/Glu: green), along with DAPI (blue) at P7, P2, and 2 months of age. White arrow indicated Ki67+ α- or β-cells. *(C)* Morphometric quantification of Ki67+ β- and α-cells shown as percentage of total number of β- or α-cells at P7, P21, and 2 months. Red and green dots indicate α- and β-cell data points, respectively. White bars represent β-cell data, while dotted bars correspond to P21 data. Error-bars show SEM. *$P<0.05$, ** $P<0.01$, using a paired, two-tailed *t*-test. Scale bar: 50 μm.

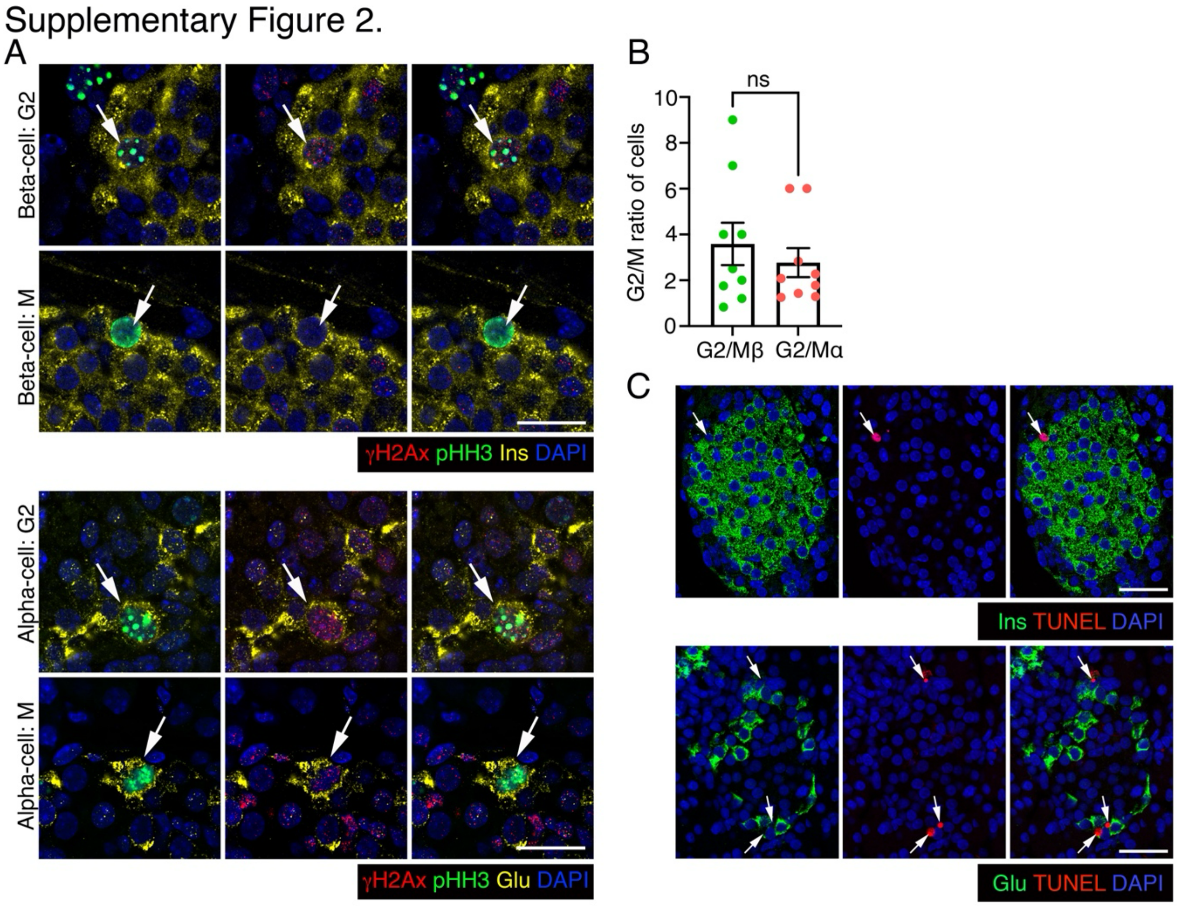


**Supplementary Figure 2 (Related to Figure 2).** *(A)* Representative Airy-scan confocal images showing ɣH2AX (red) and pHH3 (green) in β-cells (top panels) or α-cells (bottom panels), with insulin (Ins) and Glucagon (Glu) shown in yellow, along with DAPI (blue) at P7. Punctate and solid pHH3 patterns indicate the G2 and M phases, respectively. *(B)* Quantification of G2/M ratio in β- and α-cells, corresponding to Figure 2C. *(C)* Single channel images corresponding to Figure 2G, showing TUNEL (red) in β- or α-cells labeled with insulin or glucagon (Ins/Glu: green), along with DAPI (blue). White arrow indicated TUNEL+ α- or β-cells. Red and green dots indicate α- and β-cell data points, respectively. Error-bars show SEM. Ns – not significant, using a paired, two-tailed *t*-test. Scale bar: 20 µm *(A)*, 50 µm *(C)*.

Supplementary Figure 3.

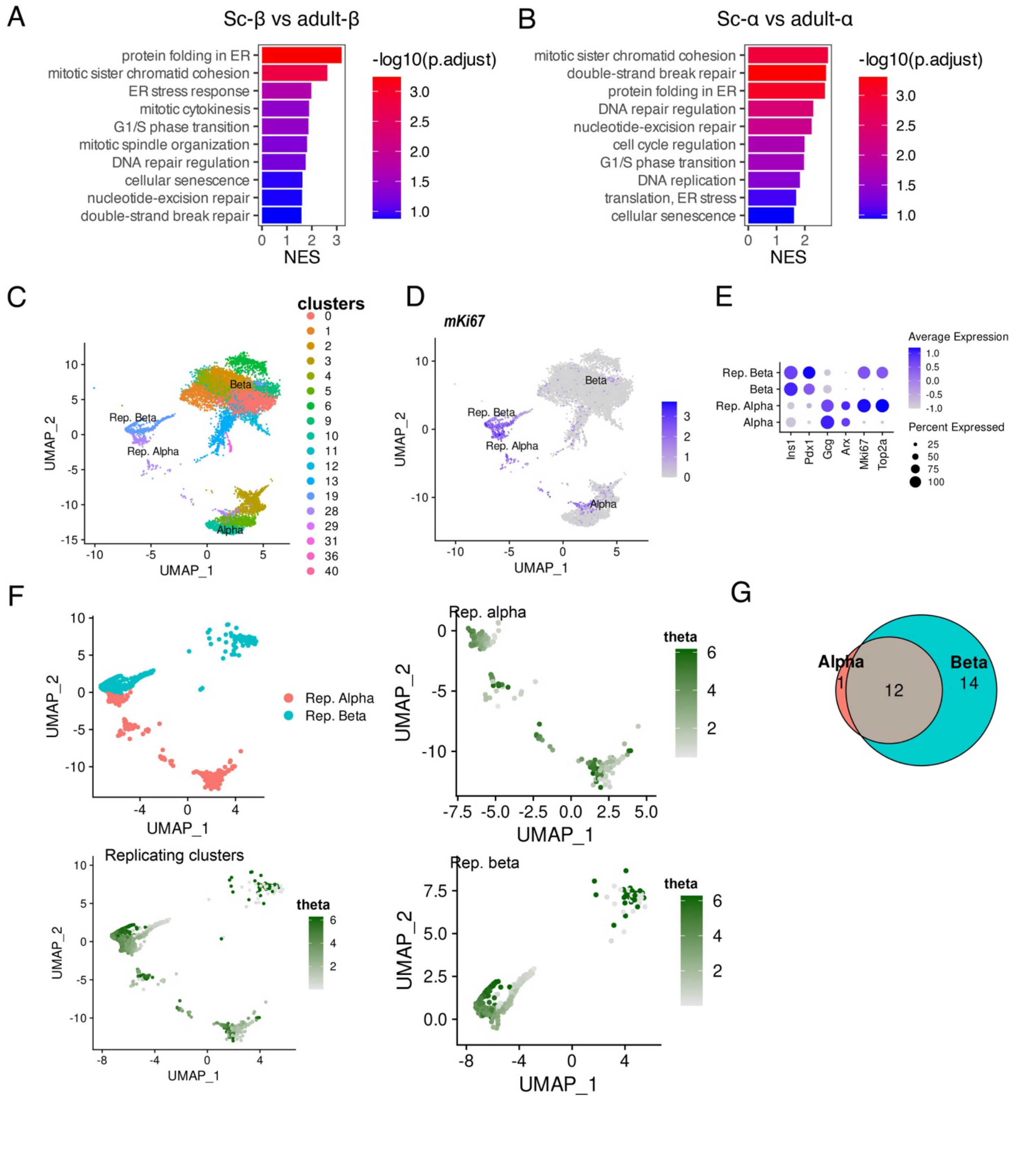


**Supplementary Figure 3 (Related to Figure 3).** *(A-B)* Top DDR and replication pathways enriched (GSEA using GO-BP, *adj.P*<0.1) in human stem-cell derived (SC-) β- *(A)* and α-cells *(B)* compared to their adult, human islet-derived counterparts(1; 2). *(C)* UMAP projection showing identified cell clusters in mouse neonatal islet scRNA-seq. *(D)* Replicative α- and β-cells are highlighted in UMAP space via their *Mki67* expression. *(E)* Dot-plot showing the expressions of cell identity and replicative marker genes for validation of replicative α- and β-cell identities. *(F)* UMAP plot of replicative α- and β-cells colored by the cell type (top left panel) and the cell cycle identifier theta (rest of the three panels) derived using Tricycle. *(G)* Venn-diagram comparing DDR processes that show significant (*P*<0.05) differences in their GSVA scores between G2 and M phases in replicative α- and β-cells.

Supplementary Figure 4.

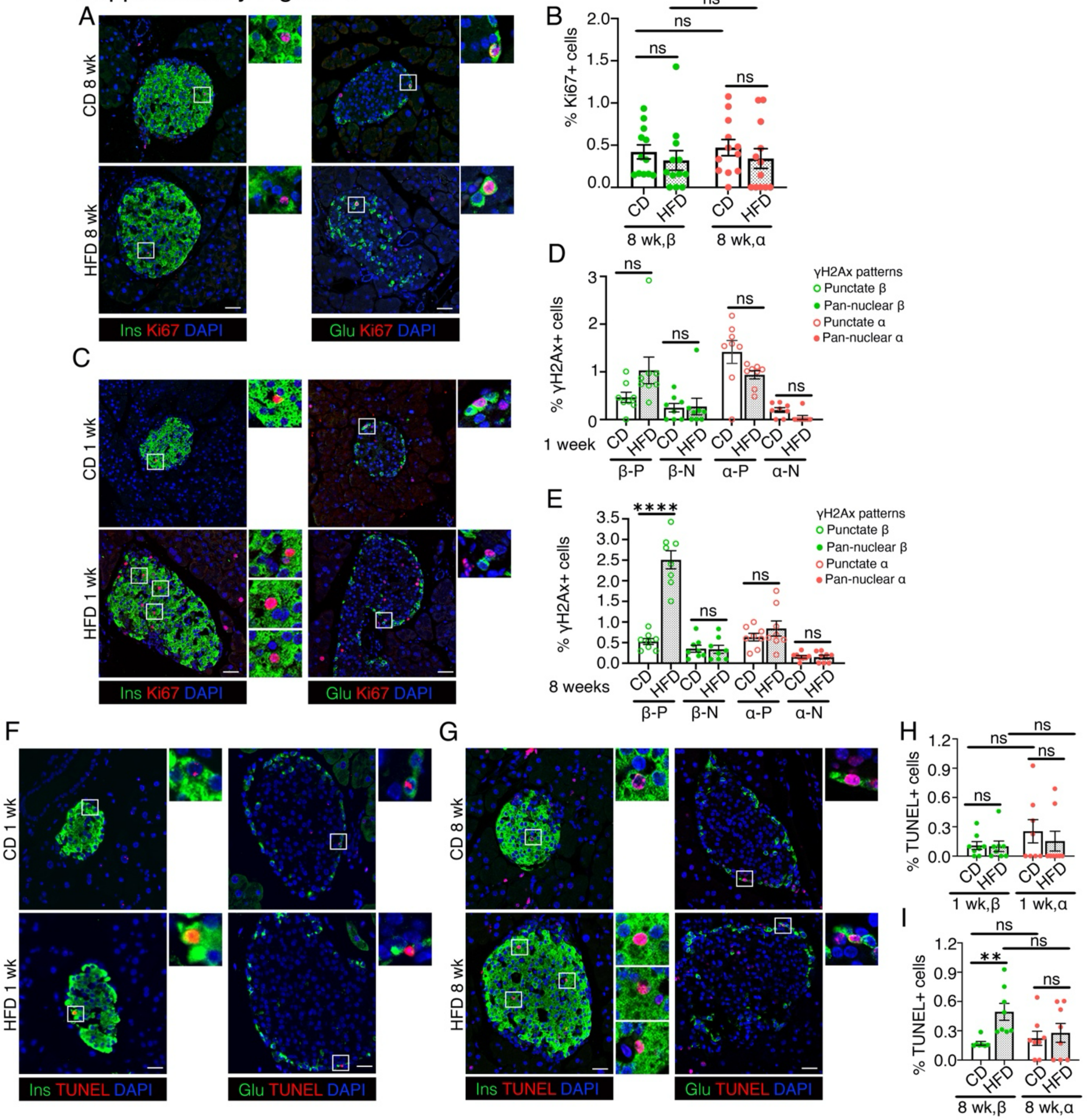


**Supplementary Figure 4 (Related to Figure 4).** *(A, B)* Representative images *(A)* and quantification *(B)* for Ki67 (red) in β-cells (left panels) or α-cells (right panels), with insulin or glucagon (green) and DAPI (blue) in pancreatic sections from mice fed with a chow diet (CD) or a high fat diet (HFD) for 8 weeks. Representative areas marked by white square corresponding to each image are presented as a 2X magnified panel next to the image. Data in *(B)* shows Ki67 quantification in β- and α-cells as percentage of total number of β- or α-cells in CD and HFD mice. *(C)* Representative images for Ki67 (red) in β-cells (left panels) or α-cells (right panels), with insulin or glucagon (green) and DAPI (blue) in pancreatic sections from CD control and 1-week HFD mice. Representative areas marked by white square corresponding to each image are presented as a 2X magnified panel next to the image. *(D, E)* Quantification of ɣH2AX+ β- and α-cells with punctate ɣH2AX+ (P) or pan-nuclear ɣH2AX+ (N) patterns shown as percentage of total number of β- or α-cells, in 1-week *(D)* and 8 weeks *(E)* HFD and CD cohorts. *(F, G)* Representative images for TUNEL (red) in β-cells (left panels) or α-cells (right panels), with insulin or glucagon (green) and DAPI (blue) in pancreatic sections from 1-week HFD *(F)* and 8 weeks HFD *(G)* cohorts with age-matched CD controls. Areas marked by white square corresponding to each image are presented as a 2X magnified panel next to the image. *(H, I)* Quantification of TUNEL+ α- or β-cells in 1-week HFD *(H)* and 8 weeks HFD *(I)* cohorts with age-matched CD controls, shown as percentage of total number of α- or β-cells. All the images and quantification data shown is obtained from wildtype C57BL/6J mice fed with an HFD or CD for 1 week (n=7 mice for *D*, *H*) or 8 weeks (n=12 mice for *B,* n=8 for *E, I*), starting at 6 weeks of age. *(B, D, E, H, I)* Red and green dots indicate α- and β-cell data points, respectively; white bars denote control CD data, dotted bars indicate HFD data. Error-bars show SEM. Ns= not significant, ** $P<0.01$, ****$P<0.001$, using 2-way ANOVA with Fisher's LSD test for *(B, D-G)*. Scale bar: 50 μm.

Supplementary Figure 5.

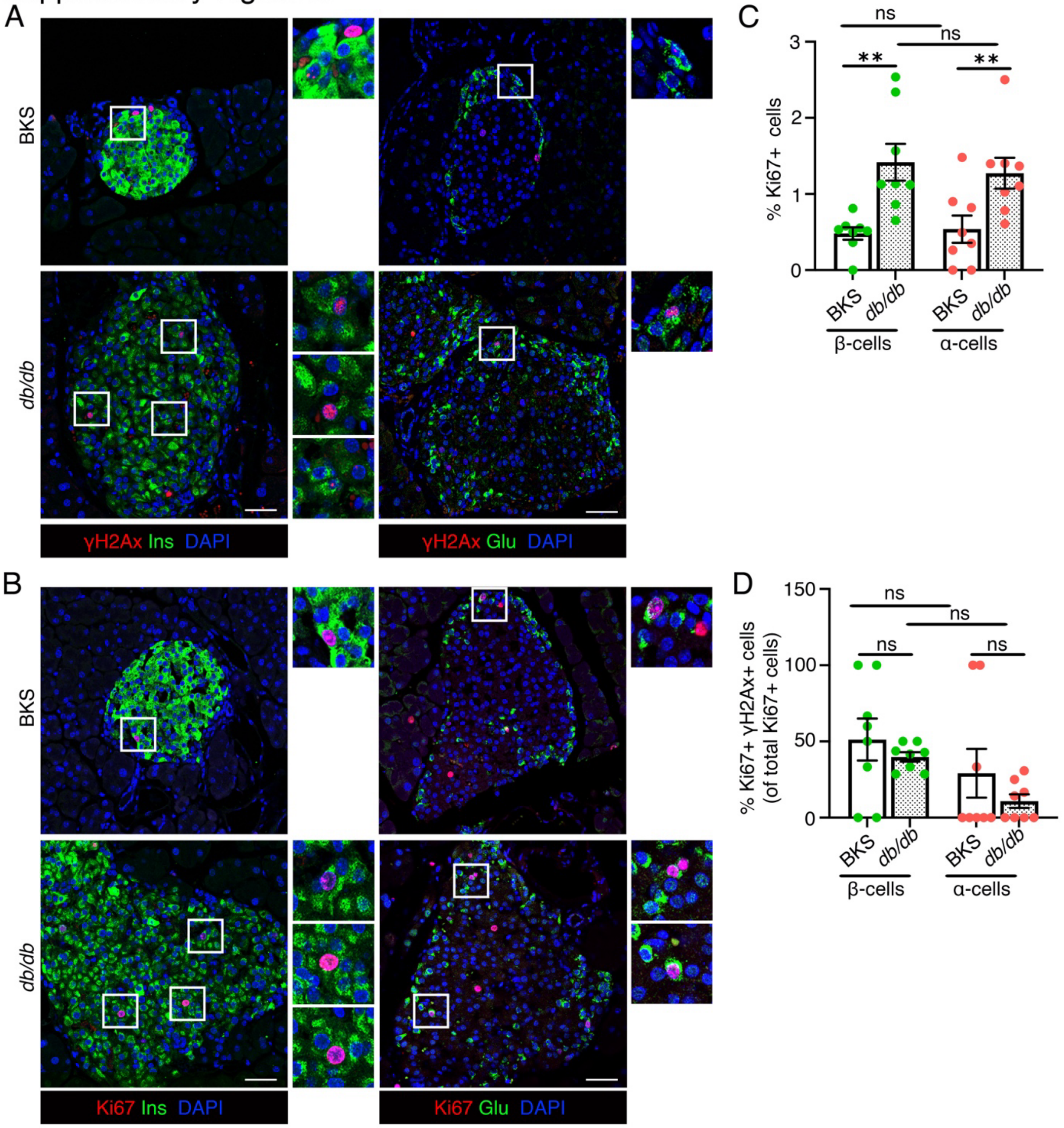


**Supplementary Figure 5 (Related to Figure 5).** *(A, B)* Representative images showing Immunostaining for ɣH2AX (A) or Ki67 (B) (red) and β- or α-cells marked by insulin or glucagon (Ins/Glu: green), along with DAPI (blue) in BKS and *db/db* pancreatic sections. *(C)* Quantification of Ki67+ β- and α-cells shown as percentage of total number of β- or α-cells in BKS and *db/db* samples. *(D)* Quantification of ɣH2AX+Ki67+ β- and α-cells shown as percentage of Ki67+ β- or α-cells in the BKS and *db/db* samples, showing fraction of the replicating endocrine cells marked by DNA damage. All the images and quantification data shown is obtained from 8 weeks old BKS and *db/db mice* (n=8/group). Red and green dots indicate α- and β-cell data points, respectively; white bars denote BKS data while dotted bars show *db/db* data. Representative areas marked by white square for each image are presented as a corresponding 2X magnification next to the image.

Error-bars show SEM. Ns= not significant, ** $P<0.01$, using 2-way ANOVA with Fisher's LSD test for *(C, D)*. Scale bar: 50 µm.

Supplementary Figure 6.

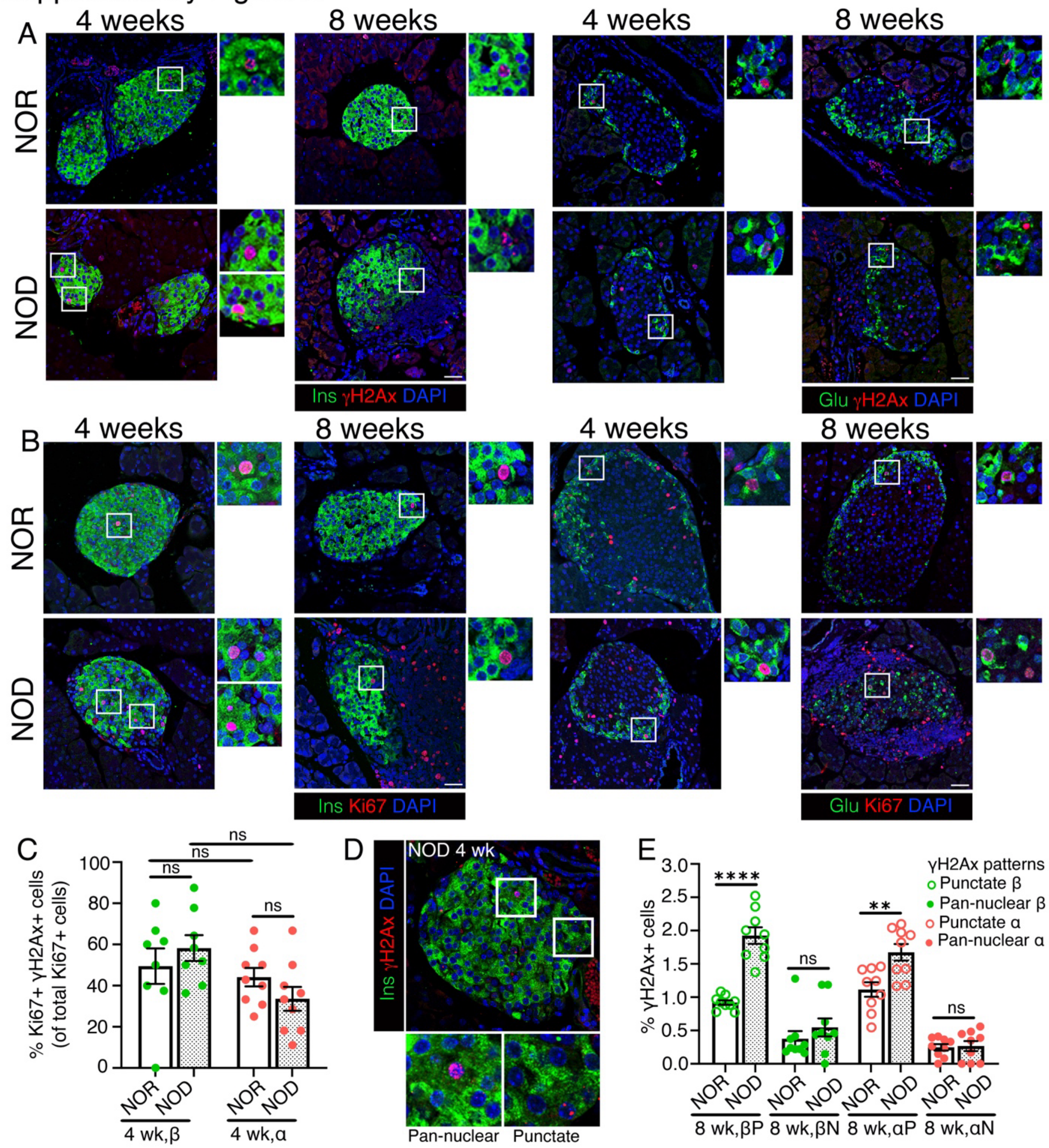


**Supplementary Figure 6 (Related to Figure 6).** *(A, B)* Representative images showing Immunostaining for ɣH2AX *(A)* or Ki67 *(B)* (red) and β- or α-cells marked by insulin or glucagon (Ins/Glu: green), along with DAPI (blue) in NOR and NOD pancreatic sections. *(C)* Quantification of ɣH2AX+Ki67+ β- and α-cells shown as percentage of Ki67+ β- or α-cells in pancreatic sections from 4 weeks old NOR and NOD mice, showing the fraction of replicating cells marked by DNA damage. *(D)* Representative immunostaining from a NOD (age = 4 weeks) pancreatic section showing ɣH2AX (red) in punctate and pan-nuclear patterns (as indicated) in β-cells (Ins: green), with DAPI in blue. *(E)* Quantification of ɣH2AX+ β- and α-cells with punctate ɣH2AX+ (P) or pan-nuclear ɣH2AX+ (N) patterns shown as percentage of total number of β- or α-cells, in 8 weeks old NOD and NOR mice. All the images and quantification data shown is obtained from 4 weeks old NOR and NOD mice (n=9/group), except data in (E), which is obtained from 8 weeks old NOD and NOR cohorts (n=9/group). Red and green dots indicate α- and β-cell data points, respectively; white bars show NOR data while dotted bars represent NOD data. Representative areas marked by white square for each image are presented as a corresponding 2X magnification next to the image. Error-bars show SEM. Ns= not significant, ** $P<0.01$, ****$P<0.001$, using 2-way ANOVA with Fisher's LSD test for *(C)* and a paired, two-tailed *t*-tests for *(E)*. Scale bar: 50 μm.

Supplementary Figure 7.

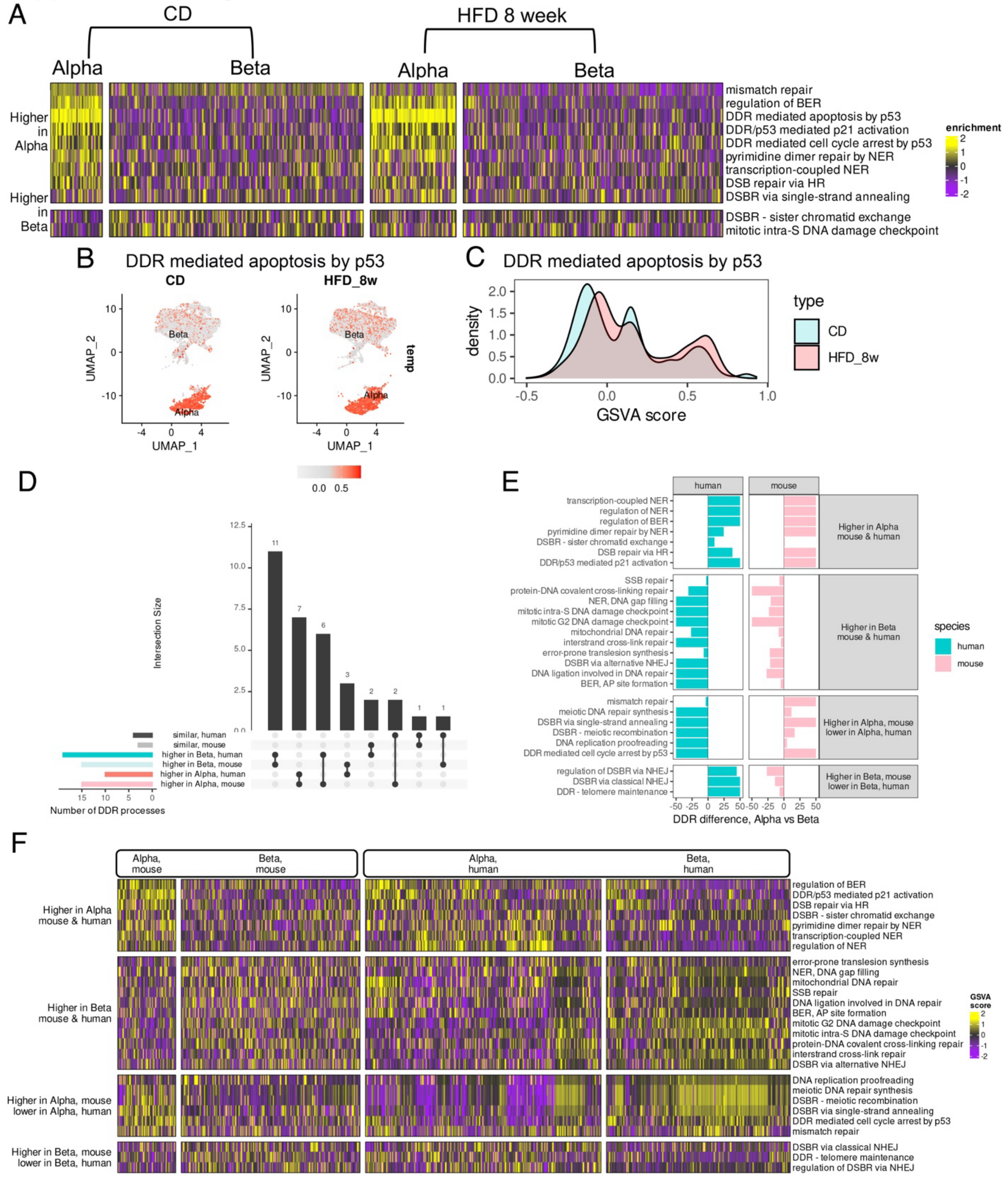


**Supplementary Figure 7 (related to Figure 7).** *(A)* Heatmap of GSVA scores of differentially regulated DDR processes between murine α- and β-cells from publicly available scRNA-seq dataset(3), compared between C57BL/6J mice fed with Control (CD) and high fat diets (HFD) for 8 weeks. *(B)* UMAP plots of α- and β-cells from CD and HFD 8-week samples, colored by the GSVA score of the DDR mediated apoptosis process. *(C)* Density distributions comparing the

GSVA scores of the DDR mediated apoptosis process between CD and HFD 8-week samples. *(D)* Upset plot comparing the DDR differences between α- and β-cells among adult mouse (chow/LFD fed adult C57BL/6J mice, (4)) and human islet samples (non-diabetic donors ages 30-50 years, HPAP)(5). The DDR processes are categorized into similar, higher in α- or higher in β-cells individually for the two species based on their GSVA score differences. *(E)* Differentially regulated DDR processes between α- and β-cells compared between mouse and human samples. The DDR processes are categorized based on similar or opposite trends between mouse and human. X-axis represents the magnitude of GSVA score difference between α- and β-cells, quantified as $-\log_{10}(P)$. The sign indicates the direction of change. Positive values indicate higher enrichment in α-cells, and negative values indicate lower in α-cells (and higher in β-cells). *(F)* Heatmap comparing the GSVA scores of differentially enriched DDR processes between α- and β-cells separately for mouse and human adult islet samples. Processes are categorized based on similar and opposing trends between the two species (mouse and human). The pathways highlighted as "Higher in α-mouse and lower in α-humans" also indicate pathways that are, by inference, also higher in human β-cells.

Supplementary Figure 8.

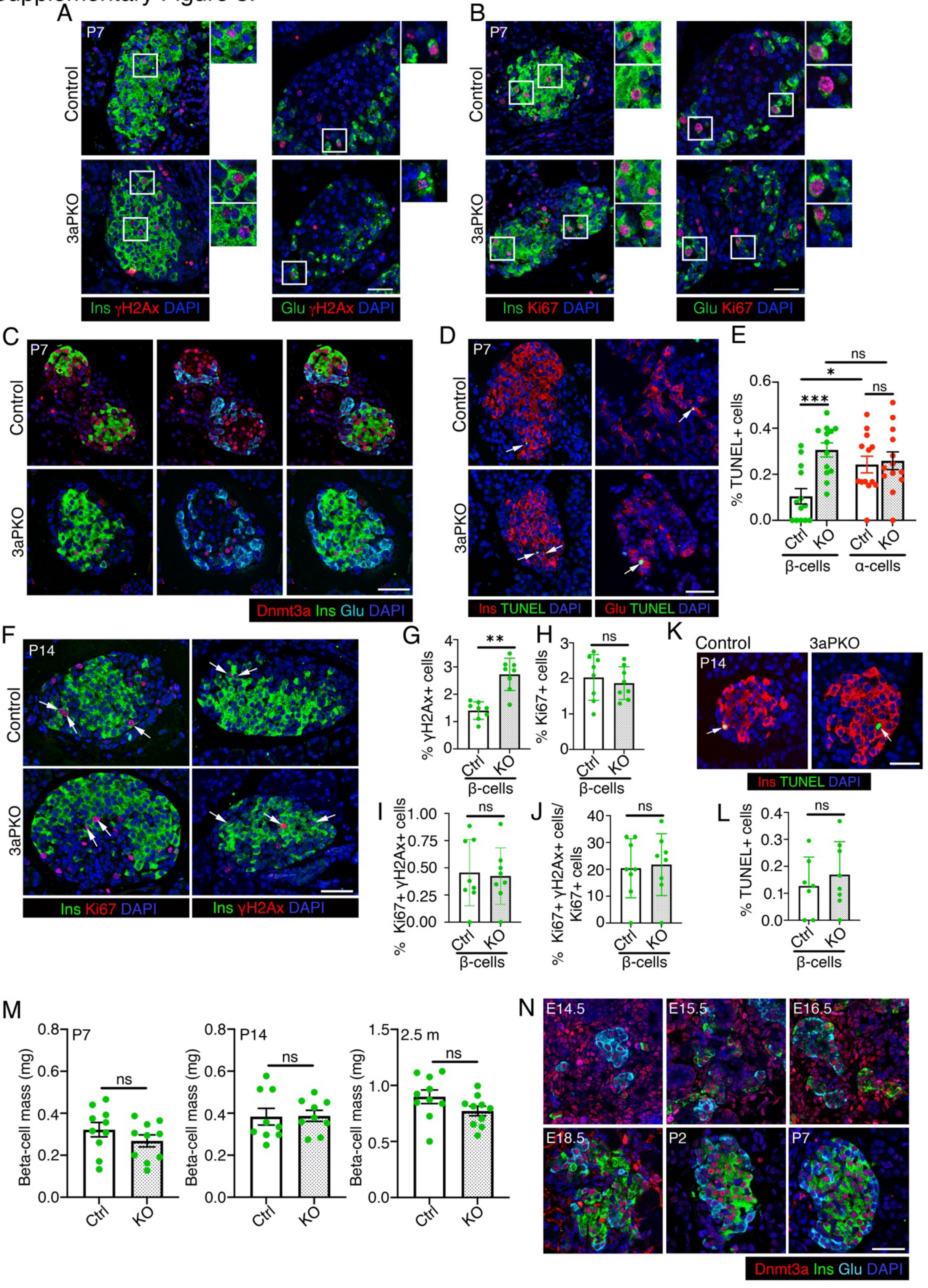

**Figure S8 (Related to Figure 8).** *(A, B)* Representative images showing Immunostaining for ɣH2AX *(A)* or Ki67 *(B)* (red) and β- or α-cells marked by insulin or glucagon (Ins/Glu: green), along with DAPI (blue) in P7 control and 3aPKO pancreatic sections. Representative areas marked by white square for each image are presented as a corresponding 2X magnification next to the image. *(C)* Immunostaining for Dnmt3a (red) along with Insulin (Ins, green) and Glucagon (Glu, cyan), along with DAPI (blue) in p7 control and 3aPKO pancreatic sections, showing loss of Dnmt3a in the KO islets. *(D, E)* Representative images *(D)* for TUNEL and quantification *(E)* in β-cells and α-cells in control and 3aPKO pups at P7. In *(D)* TUNEL (green) is shown with Ins/Glu(red) and DAPI (blue), arrows highlighting α- or β-cells marked by TUNEL. The graph in *(E)* shows TUNEL+ β- and α-cells shown as percentage of total number of β- or α-cells. *(F)* Representative immunostaining images for ɣH2AX and Ki67 (red) in β-cells, with insulin (Ins) shown in green, along with DAPI (blue) in control and 3aPKO mice at P14. Arrows indicate ɣH2AX+ or Ki67+ β-cells. *(G)* Quantification of ɣH2AX+ β-cells shown as percentage of total number of β-cells in P14 control and 3aPKO mice. *(H)* Quantification of Ki67+ β-cells shown as percentage of total number of β-cells in 3aPKO and control mice at P14. *(I)* Quantification of Ki67+ɣH2AX+ β-cells in 3aPKO and control P14 mice, shown as percentage of total β-cells, indicating the total pool of replicating cells with DNA damage. *(J)* Quantification of ɣH2AX+Ki67+ β-cells shown as percentage of Ki67+ β-cells in P14 control and 3aPKO samples, showing fraction of the replicating cells harboring DNA damage. *(K, L)* Representative images *(K)* for TUNEL and quantification *(L)* in β-cells in control and 3aPKO pups at P14. *(M)* Β-cell mass in control and 3aPKO mice at P7, P14, and 2.5 months of age. *(N)* Immunostaining for Dnmt3a (red) along with Insulin (Ins, green) and Glucagon (Glu, cyan), along with DAPI (blue) in wildtype C57BL/6J mouse pancreatic samples taken from indicated embryonic and neonatal stages. All the images and quantification data shown is obtained from control and 3aPKO at P7 (n=14 A-C, n=13 D-E, n=10 M) p14 (n=8 F-J, n=7 K-L, n=9 M) and n=5 for panel N. Red and green dots indicate α- and β-cell data points, respectively; white bars denote control data, dotted bars indicate 3aPKO data. Error-bars show SEM. Ns= not significant, **P<0.05,* ** *P*<0.01, ****P*<0.005, ****P<0.001, using 2-way ANOVA with Fisher's LSD test for *(E)*, and unpaired, two-tailed *t*-test for *(G-J, L, M)*. Scale bar: 50 μm.

**Supplementary Table 1**

**Antibodies used for immunofluorescence analyses (primary and secondary)**

| **Antibody** | **Dilution** | **Source and Catalog Number** |
|---|---|---|
| Mouse monoclonal anti-Glucagon | 1:1000 | Sigma-Aldrich G2654-.2ML |
| Rabbit polyclonal anti-Glucagon | 1:500 | Immunostar 20076 |
| Guinea polyclonal pig anti-Glucagon | 1:2000 | Progen 16032 |
| Guinea polyclonal pig anti-Insulin | 1:20,000 | Biosynth 20-1P35 |
| Rabbit polyclonal anti-insulin | 1:20,000 | Abcam ab181547 |
| Mouse monoclonal anti-Insulin | 1:2,000 | Millipore-Sigma I2018-100UL |
| Mouse monoclonal anti-Ki67 | 1:100 | BD Biosciences 550609 |
| Mouse monoclonal anti-pHH3 | 1:2,000 | Abcam ab14955 |
| Rabbit polyclonal anti-pHH3 | 1:200 | Millipore-Sigma 06-570 |
| Rabbit monoclonal anti-ɣH2Ax | 1:500 | Cell Signaling 9718 |
| Mouse monoclonal anti-ɣH2Ax | 1:1,000 | Abcam 2118009 |
| Mouse anti-Dnmt3a | 1:200 | Novus Biologicals NB120-13888 |
| Cy™3 AffiniPure Donkey Anti-Rabbit IgG (H+L) | 1:2000 | Jackson ImmunoResearch 711-165-152 |
| Cy™3 AffiniPure F'(ab)2 fragment Donkey Anti-Mouse IgG (H+L) | 1:500 | Jackson ImmunoResearch 715-166-151 |
| Alexa Fluor® 488 AffiniPure Donkey Anti-Guinea Pig IgG (H+L) | 1:1,000 | Jackson ImmunoResearch 706-545-148 |
| Alexa Fluor® 488 AffiniPure Donkey Anti-Rabbit IgG (H+L) | 1:1,000 | Jackson ImmunoResearch 711-545-152 |
| Alexa Fluor® 488 AffiniPure Donkey Anti-Mouse IgG (H+L) | 1:500 | Jackson ImmunoResearch 715-545-150 |
| Alexa Fluor® 647 AffiniPure Donkey Anti-Guinea Pig IgG (H+L) | 1:500 | Jackson ImmunoResearch 706-605-148 |
| Alexa Fluor® 647 AffiniPure Donkey Anti-Rabbit IgG (H+L) | 1:500 | Jackson ImmunoResearch 711-605-152 |
| Alexa Fluor® 647 AffiniPure F'(ab)2 fragment Donkey Anti-Mouse IgG (H+L) | 1:500 | Jackson ImmunoResearch 715-606-150, |

**Supplementary Table 2.** Cell type specific markers

| **Alpha** | **Beta** | **Delta** | **PP** | **Ductal** | **Activated stellate** | **Quiescent stellate** | **Endothelial** | **Macrophage** | **T cell** | **B cell** |
|---|---|---|---|---|---|---|---|---|---|---|
| *Gcg*<br>*Ttr*<br>*Irx2*<br>*Arx*<br>*Mafb* | *Ins1*<br>*Ins2*<br>*Mafa*<br>*Nkx6-1*<br>*Slc2a2*<br>*G6pc2* | *Sst*<br>*Ghsr*<br>*Rbp4*<br>*Hhex* | *Ppy* | *Krt19*<br>*Hnf1b*<br>*Sox9* | *Col1a1*<br>*Timp1*<br>*Pdgfra* | *Gja4*<br>*Ednrb*<br>*Col4a1* | *Plvap*<br>*Esm1* | *Adgre1*<br>*Lyz2* | *Trbc2*<br>*Cd3g* | *Cd19*<br>*Igkc*<br>*Ighm* |

### Supplementary Methods

#### *Immunohistochemical staining*

Briefly, FFPE pancreatic sections were deparaffinized using toluene and gradients of ethanol and rehydrated, followed by heat-mediated antigen retrieval using a citrate buffer-based antigen unmasking solution (Vector Labs). Slides were incubated in 0.4% triton-TBS for 30 minutes, followed by blocking for one hour in 0.2% Tween-TBS with 3% BSA. Slides were then incubated with a cocktail of primary antibodies prepared in blocking buffer at concentrations indicated in Supplementary Table 1 O/N at 4°C. Slides were washed next day in TBST (TBS + 0.2% Tween20) twice (15' per wash) followed by a single wash with TBS (15' per wash) and then mounted in an antifade mounting medium containing DAPI (Vector Labs).

#### *TUNEL staining*

Cell death was measured by the **T**erminal Deoxynucleotidyl Transferase (TdT)-mediated d**U**TP **N**ick-**E**nd **L**abeling (TUNEL) assay, performed using the DeadEnd™ Fluorometric TUNEL System kit (Promega) according to the manufacturer's instructions. Briefly, FFPE pancreatic sections were deparaffinized, rehydrated, and fixed with 4% formaldehyde for 15 minutes. After washing with PBS, sections were treated with proteinase K (20µg/mL) for 8 minutes, washed again with PBS, fixed, and washed once more before proceeding with the TdT reaction for 90 minutes. After the reaction and incubation with SSC, slides were blocked and stained for insulin or glucagon.

#### *Bulk RNA-seq analysis*

Paired-end RNA-Seq reads were trimmed to remove sequencing adapters using Trimmomatic (version 0.38(6)) with parameters LEADING:3, TRAILING:3, SLIDINGWINDOW:4:15, and MINLEN:36. PolyA tails were trimmed using FASTP(7) with default settings. The resulting processed reads were aligned to the mouse reference genome (mm10) using STAR (version 2.6.0a(8)) with options --twopassMode Basic and --outSAMstrandField intronMotif. Gene-level read counts were generated using HTSeq (version 0.11.1(9)) with default parameters to produce the count matrix. Differential expression analysis was conducted by adjusting read counts to normalized expression values using TMM normalization method in edgeR(10). Briefly, for between phenotype comparison, general linear models were applied to identify DEGs using TMM normalization expression level as a depending variable, and phenotype as an independent variable. Genes with an FDR-adjusted *p*-value less than 0.05 and with a fold change (FC) greater than 1.5 or less than 0.5 were considered as significant up- and down-regulated genes, respectively. Gene set enrichment analysis (GSEA) was conducted using the Gene Ontology Biological Process (GO:BP) dataset(11), employing the R package ClusterProfiler V4.0(12). Consistent with recommendations in the official GSEA user guide, a false discovery rate (FDR) threshold of 25% (FDR < 0.25) was applied to determine significantly enriched gene sets, a standard criterion for exploratory analyses. In addition to our in-house neonatal and adult bulk RNA-seq datasets, we used bulk RNA-seq data on sorted adult mouse α- and β-cells generated by Benner and colleagues(13).

***Single cell RNA-seq analysis***

Raw fastq reads from the in-house neonatal murine samples were aligned to the mouse genome (mm10) followed by gene level quantification in CellRanger V7.0 (14). The data were cleaned by removing doublets in DoubletFinder V2.0(15), followed by ambient RNA correction using SoupX V1.6.2(16). Next, low quality cells were removed based on low cytoplasmic RNA or high mitochondrial RNA followed by normalization using SCtransform(17). The in-house samples, along with the adult murine samples from the published datasets were integrated using Harmony V1.2.4(18), followed by UMAP projection and clustering (Leiden) in Seurat V4.0(17; 19-22). The cell type for each cluster was determined based on the expressions of known marker genes(23; 24) (Supplementary Table 2). The identities of the replicative α- and β-cell clusters were determined based on the expressions of *Mki67* and *Top2a*. Normalized and integrated single-cell data from human islets was obtained from HPAP(5; 25-27) by downloading from the PANC-DB website and analyzed in Seurat V4.0(17; 19-22).

Enriched DDR pathways in neonatal mouse α- and β-cells were obtained using Geneset enrichment analysis (GSEA) on the gene-wise log fold changes between neonatal (in-house) and adult (islets from C57BL/6J mice fed with LFD chow diet)(4) samples as implemented in the R package ClusterProfiler V4.0(12). A list of 160 DDR pathways and associated genesets were obtained from the GO database using the R package GO.db(11; 28). To avoid including redundant pathways, the GO-BP hierarchy was analyzed to retain only the terminal pathways from each branch, giving a total of 35 non-redundant pathways. Genset variation analysis (GSVA) of the murine and human samples (HPAP) were performed on the normalized counts matrices using a gaussian kernel as implemented in the R package GSVA, employing the non-redundant DDR process genesets(29). Differentially regulated pathways between two conditions were identified based on pvalues obtained from Student's *t*-test. Cell cycle phases for the replicative α- and β-cells were identified separately for each sample using the program Tricycle, following the instructions posted on the Bioconductor website(30; 31). Visualization of gene expression profiles in UMAP, heatmaps, barplots and upset plots were created using the packages Seurat(17; 19-22), complexHeatmap(32) ggplot(33), and UpsetR(34) respectively.

***Summary of publicly available datasets used***

We used scRNA-seq datasets from C57BL/6J mice fed with low fat/chow diet (LFD/CD) or HFD for 1-week (4)(GEO accession ID: GSE162512) and 8-weeks (GEO accession ID: GSE203376)(3), and bulk RNA-seq of sorted adult C57BL6/J mouse α- and β-cells(13). For human stem-cell derived (SC) α- and β-cells, we used the dataset generated by Balboa and colleagues (GEO accession ID: GSE167880)(1), compared to adult human islet scRNA-seq data used by them(GEO accession ID: GSE114297)(2) For data in (Supplementary Figs 3A, B), we used the combined data as presented in the Balboa study. scRNA-seq data for non-diabetic and T2D adult human islets (30-50 years of age) was downloaded from PANC-DB, an HPAP resource(5; 25-27) and used for comparison of ND and T2D human α- and β-cells as well as compared with (3) to derive conserved and distinct DDR pathways between mouse and human α- and β-cells. This dataset is available at https://hpap.pmacs.upenn.edu/.
